# Cryptic functional diversity in ant tandem runs

**DOI:** 10.1101/2022.08.28.505613

**Authors:** Nobuaki Mizumoto, Yasunari Tanaka, Gabriele Valentini, Thomas O. Richardson, Sumana Annagiri, Stephen C Pratt, Hiroyuki Shimoji

**Affiliations:** Okinawa Institute of Science & Technology Graduate University, Onna-son, Okinawa, 904-0495, Japan; School of Biological and Environmental Sciences, Kwansei Gakuin University, Sanda, Hyogo, 669-1337, Japan; IRIDIA, Université Libre de Bruxelles, Brussels, Belgium; School of Biological Sciences, University of Bristol, UK; Behaviour and Ecology Lab, Department of Biological Sciences, Indian Institute of Science Education and Research Kolkata, West Bengal 741246, India; School of Life Sciences, Arizona State University, Tempe, AZ 85287, USA

**Author notes:** These authors contributed to the work equally (Nobuaki Mizumoto, Yasunari Tanaka).

## Abstract

Ants have evolved diverse recruitment methods to guide colony members to valuable resources, such as food or nest sites. One of these methods, tandem running, consists of an informed leader directly guiding a naive follower every step of the way from nest to resource. Although this behavior appears superficially similar in the different ant taxa in which it has independently evolved, this similarity could conceal underlying functional and mechanistic differences. Here we present a combined network and information-theoretic analysis, which reveals fundamental differences in the tandem recruitment between two distantly related ant genera, *Temnothorax* and *Diacamma. Temnothorax* uses tandem running to recruit additional recruiters, whereas *Diacamma* uses it principally to move the passive majority of the colony, a task that *Temnothorax* accomplishes with a different behavior, social carrying. Accordingly, the structure of the tandem run recruitment networks of *Diacamma* was different from those of *Temnothorax*, with *Diacamma* networks more closely resembling the social carrying networks of *Temnothorax*. Furthermore, an information-theoretic analysis of the spatial trajectories of leaders and followers revealed that *Diacamma* tandem runs lack bidirectional information flow, the signature of route-learning in *Temnothorax* tandem runs. These results suggest that *Diacamma* uses tandem runs not to share information, but to transport nestmates. By quantifying the cryptic diversity of communication behavior, this study increases the resolution of our understanding of animal societies.

## Introduction

Social insects cooperatively accomplish an impressive array of collective tasks, including construction of complex nest structures, flexible division of labor, and mass recruitment of colony members (Wilson, 1971). These collective behaviors are underpinned by their communication systems, which allow social insect colonies to quickly disseminate valuable information among colony members (Leonhardt et al., 2016). The social insects have evolved a diversity of communication systems, ranging from direct one-to-one interactions (Meurville and LeBoeuf, 2021; Tsuji et al., 1999) to indirect interactions through chemical signals deposited in the shared environment (Camazine et al., 2001; Theraulaz and Bonabeau, 1995). The diversification of communication systems has been accompanied by instances of remarkable evolutionary convergence. For example, *Lasius* ants and *Nasutitermes* termites produce similar sponge-like nest structures (Perna and Theraulaz, 2017), and the iconic mass raids of new and old-world army ants result from independent evolutionary origins (Borowiec, 2019). The convergent evolution of behavior is often considered an outcome of adaptation to the same evolutionary pressure (Gallant and O’Connell, 2020). However, convergence can also occur with differentiated functions or through distinct evolutionary contexts (Losos, 2011). Thus, superficially similar social behaviors may conceal substantial differences in their function and mechanisms for social interactions.

Tandem recruitment by ants is achieved by a unique communication technique. This behavior is used by an informed ant to share the location of a valuable resource by directly leading a naïve follower to it (Moglich et al., 1974). It has evolved independently multiple times (Glaser and Grüter, 2022; Reeves and Moreau, 2019), but it is especially well-documented in the genus *Temnothorax* (Möglich, 1978), where it is used both for foraging and colony emigration. In these ants, a successful tandem run is achieved through bidirectional feedback between partners (Franks and Richardson, 2006). This feedback allows the follower to regulate the movement speed of the pair, so that the follower can learn the travel route demonstrated by the leader (Sasaki et al., 2020; Valentini et al., 2020b). In other ant lineages where tandem running has arisen (Fig. 1), it is also used to guide colony members to a goal (Kaur et al., 2017; Schultheiss et al., 2015; Silva et al., 2021). However, more detailed investigation is required to determine whether tandem communication is functionally and mechanistically equivalent across different evolutionary origins.

**Figure 1.**
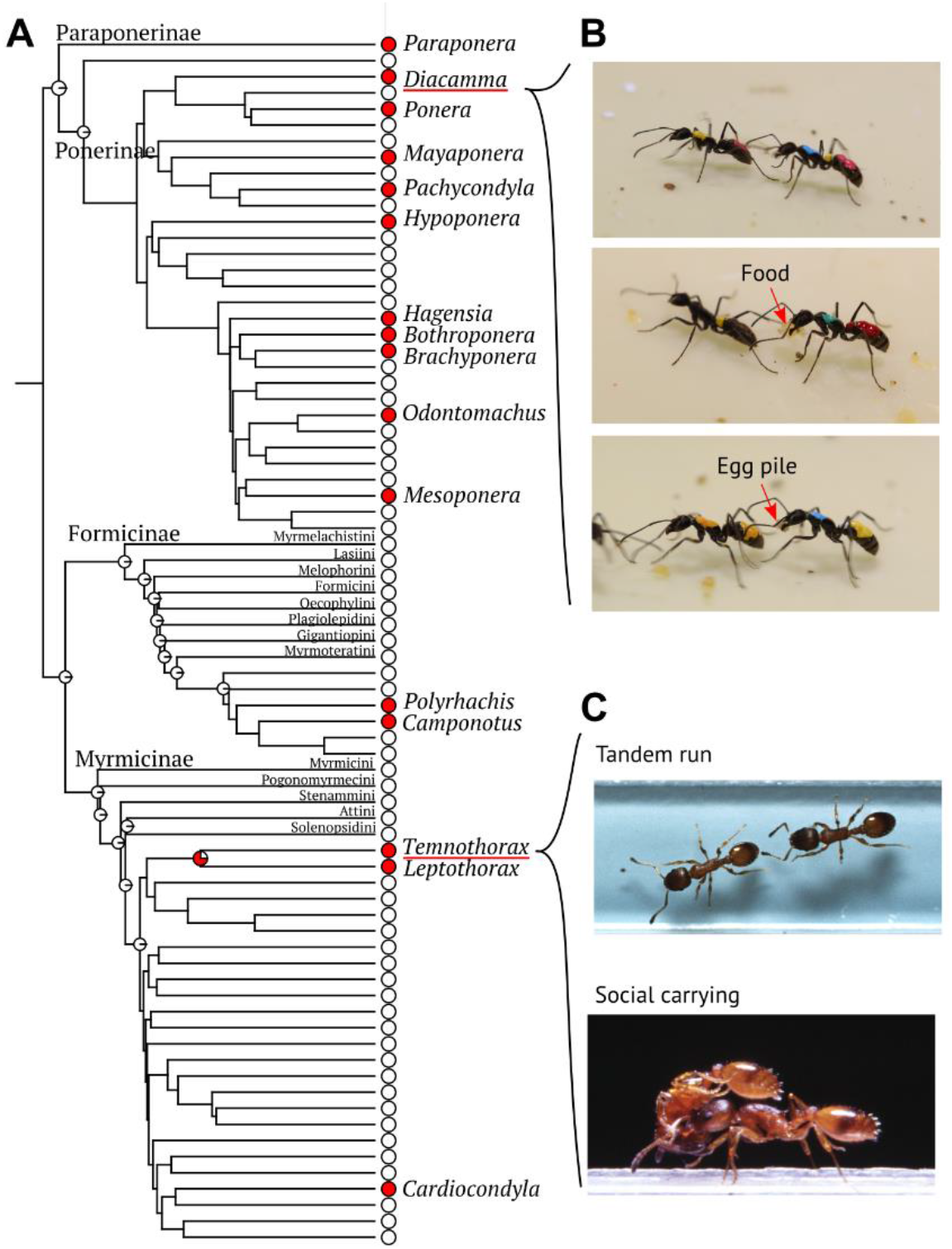
Evolution of ant tandem runs and studied species. (A) Ant phylogeny focusing on the evolution of tandem runs, simplified from (Matthew P. Nelsen et al., 2018). Red circles at tips indicate the presence of the tandem run. Node pie charts represent the probability of the estimated ancestral states of tandem run, inferred with maximum likelihood estimation methods (see Supplementary text). Estimated ancestral states are shown for roots of each tribe or where the tandem run was present with a probability larger than 70%. The full genus-level phylogeny used for ancestral state reconstruction is shown in Fig S1. Genera containing studied species are underlined in red. Tandem running evolved at least 15 times independently in ants. (B) Tandem running of *Diacamma* cf *indicum*. Followers can carry food or brood while traveling. (C) Tandem running and social carrying in *Temnothorax albipennis*.

Comparison of tandem runs by *Temnothorax* and *Diacamma* ants is a potentially revealing example of convergent evolution. Both genera use tandem runs during colony emigration, but they do so in a different manner (Fig. 1). In *Temnothorax*, tandem runs mainly occur at the initial stage of the emigration. A minority of specialized leaders recruit specialized followers to assess potential new nest sites. Once these ants decide on a site, they switch from tandem runs to social carrying, which they use to transport the rest of their colony members, one at a time, to the new nest (Möglich, 1978). Social carrying is not thought to allow the transportees to learn the emigration route, but its greater speed makes it more suitable for moving large numbers of ants that will not need to retrace the route (Pratt et al., 2005). On the other hand, *Diacamma* uses only tandem runs throughout the colony emigration (Fukumoto and Abe, 1983). All colony members move between the old and new nest sites either as leaders or followers of tandem runs. This implies that the tandem run of *Diacamma* may be functionally more similar to *Temnothorax* social carrying than to *Temnothorax* tandem runs: a means of moving the bulk of the colony rather than a means of recruiting additional recruiters.

If tandem running fulfills different functions in different species, this is reflected in the information flows between leaders and followers. For example, a previous study distinguished similar tandem running behaviors between *Temnothorax* ants and termites (Valentini et al., 2020b). In *Temnothorax* ants, tandem running allows followers to learn the route. Followers do not passively follow their leader, but frequently break contact to gather spatial information. Both the leader and the follower alternately influence their partner’s behavior: at a short distance, the leader determines the course of tandem movements, while at a long distance, the follower determines the leader’s speed (Franks and Richardson, 2006; Valentini et al., 2020b), and this two-way feedback is a consequence of teaching route information to the follower. On the other hand, in termites, tandem running is used by a mating pair and functions to keep them together as they perform a random search for nest sites: there is no need for route-learning. Thus, their interactions lack bidirectional feedback, i.e., termite followers just follow their leaders (Valentini et al., 2020b). *Diacamma* tandem runs may similarly lack route-learning protocols used by *Temnothorax*, if they primarily play a simple nestmate-moving function like social carrying. If so, then one can expect interindividual interactions in a *Diacamma* tandem pair to resemble those of termites.

Here we study the recruitment dynamics and movement coordination mechanisms of both *Temnothorax* and *Diacamma*, to reveal that tandem running in these two species is functionally and mechanistically distinct. For comparative analysis, we employed data on colony emigration and tandem runs in five different ant (sub)species: *D. indicum, D*. cf *indicum* (from Japan), *T. albipennis, T. nylanderi*, and *T. rugatulus*. We additionally used data on tandem runs in two termite species as an out-group contrast. Using these datasets, we explore whether *Diacamma* tandem running is functionally more similar to tandem running or social carrying in *Temnothorax*. First, we tested for between-species differences in the probabilities of engaging or switching between leading, following, and carrying roles. Then, we compared the recruitment network structures among species. Finally, we measured the direction of the information flow between leaders and followers within tandem runs.

## Results

### Species differences in the functions of tandem runs

During colony emigration in *Diacamma*, most individuals participated in tandem runs (Mean ± SD; *D*. cf *indicum*: 86.2 ± 8.5%; *D. indicum*: 76.0 ± 12.0%). In contrast, during emigration in *Temnothorax*, only a minority of workers engaged in tandem runs (*T. rugatulus*: 18.8 ± 9.9%). Participation in tandem running was significantly lower in *Temnothorax* than in *Diacamma*, both for leaders and followers (GLMM: *P* < 0.001, Fig 2A). *Temnothorax* instead showed a high prevalence of social carrying (*T. rugatulus*: 83.5 ± 6.0%). Indeed, the proportion *Temnothorax* workers that carried nestmates was similar to the proportion of *Diacamma* workers that led tandem runs in *D*. cf *indicum* (GLMM; vs *D*. cf *indicum*: χ^2^_1_ = 1.77, *P* = 0.18, vs *D. indicum*: χ^2^_1_ = 8.27, *P* = 0.004, Fig 2A). Similarly, the proportion of the *Diacamma* followers were similar to *Temnothorax* workers that was carried (GLMM; vs *D*. cf *indicum*: χ^2^_1_ = 0.008, *P* = 0.93, vs *D. indicum*: χ^2^_1_ = 1.54, *P* = 0.22, Fig 2A). Thus, in terms of the proportion of individuals involved, tandem running in *Diacamma* is more similar to social carrying in *Temnothorax*, rather than *Temnothorax* tandem running.

**Figure 2.**
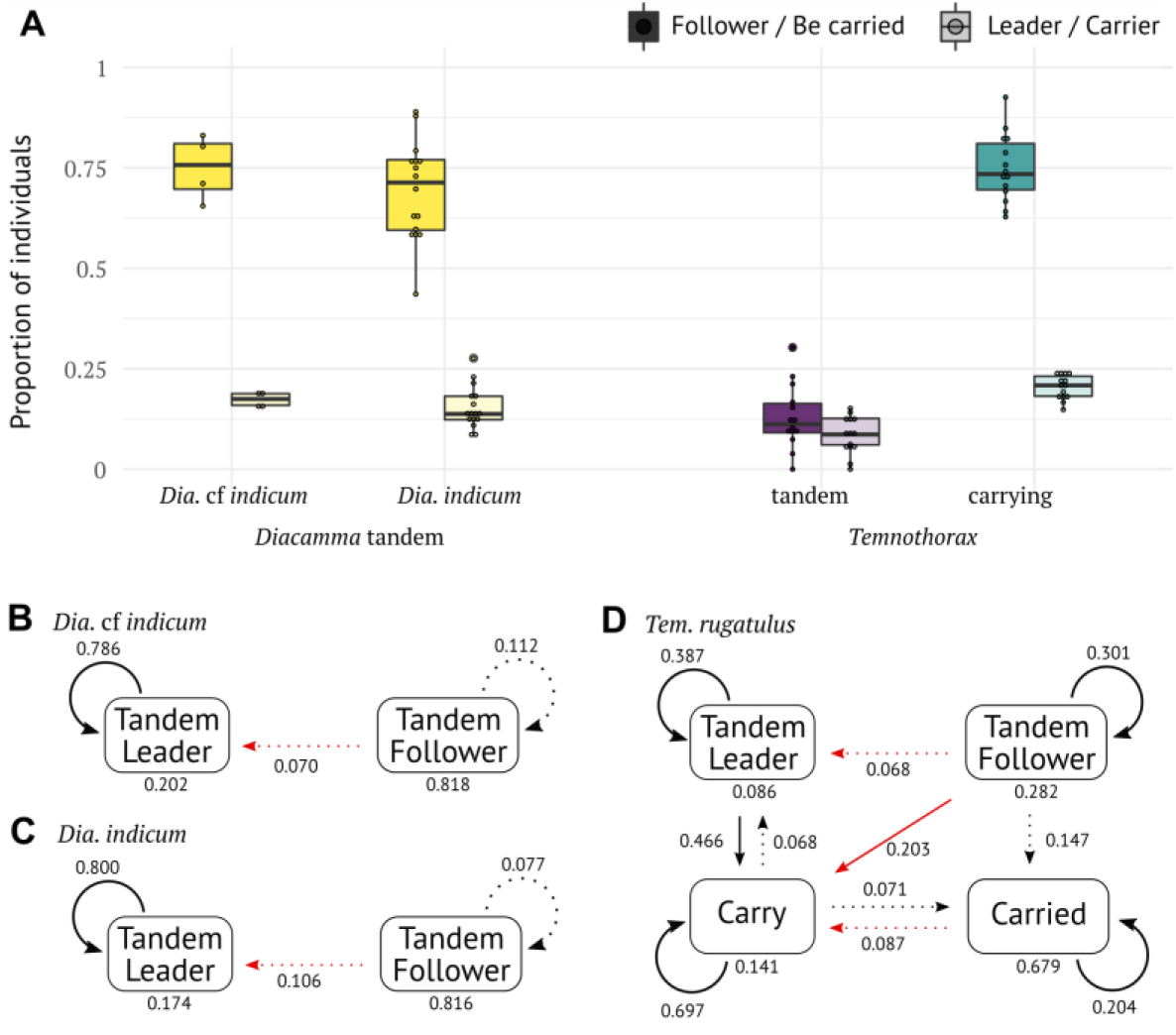
Worker participation in nestmate recruitment during colony emigration, compared between *Temnothorax* and *Diacamma*. (A) Proportion of workers involved in recruitment (tandem runs and social carrying) either as leaders/carriers or followers/carried. (B-D) Transition diagram of recruitment tasks during emigration in (B) *D*. cf *indicum*, (C) *D. indicum*, and (D) *T. rugatulus*. The results of *T. nylanderi* and *T. albipennis* are in Fig. S2. Recursive arrows indicate repetition of the same task. Numbers next to each arrow give the associated transition probability; numbers below each box indicate the probability of concluding the emigration after the task. Dashed lines show transitions with probability < 0.2; transitions with probability < 0.05 are omitted. Red lines indicate transition from being a recruit (tandem follower or being carried) to recruiting (tandem leader or carrier) roles, suggesting successful recruitment of new recruiters.

In *Temnothorax*, tandem runs function to recruit new recruiters; that is, tandem followers should go on to become tandem leaders or carriers (Pratt et al., 2005). We looked for evidence of this by measuring the transition probabilities between different recruitment roles. In *Diacamma*, we found that tandem followers rarely went on to become tandem leaders (*D*. cf *indicum*: 7%; *D. indicum*: 10.6%; Fig. 2BC). This contrasted with *T. rugatulus*, where 27.1% of tandem followers went on to become recruiters (Fig. 2D), a significantly higher proportion than in *D*. cf *indicum* (Fisher’s exact test: *P* < 0.001) or *D. indicum* (Fisher’s exact test: *P* < 0.001). Note that tandem runs of *T. nylanderi*, but not *T. albipennis*, showed a similar pattern to *T. rugatulus*, although information on social carrying was unavailable in these species (Fig. S2). On the other hand, *T. rugatulus* ants that were transported by social carrying showed a similar pattern to *Diacamma* tandem followers. Only 8.7% of carried workers became recruiters (Fig. 2D), which was not significantly different from *D*. cf *indicum* (Fisher’s exact test: *P* = 0.199) or *D. indicum* (Fisher’s exact test: *P* = 0.295).

These statistical differences between the recruitment dynamics are readily apparent in the structure of the recruitment networks during a given emigration (Fig. 3A-C, Fig. S8). The out-degree and in-degree distributions of the tandem networks differ between *Diacamma* and *Temnothorax* (KS test; in-degree: *D* = 0.26, *P* < 0.001; out-degree: *D* = 0.25, *P* < 0.001; Fig. 3DE). In *Diacamma* tandems, the recruitment networks were composed of multiple star-like structures, each with a leader as a central node and its followers as leaves (Fig. 3A). Most individuals participate in only one tandem run, as a follower, and have just one incoming edge and no outgoing edges (Fig. 3DE). In *Temnothorax* tandems, on the other hand, followers remained involved in recruitment, either by becoming leaders or by being led multiple times by different leaders during emigration (Fig. 3B). Thus, many individuals had outgoing edges with multiple incoming edges or without incoming edges (Fig. 3DE), indicating the presence of specialized leaders and specialized followers (Richardson et al., 2021, 2018; Valentini et al., 2020a). The proportion of passive workers with one incoming edge and no outgoing edge was significantly lower in *Temnothorax* tandem than *Diacamma* tandem (LMM; Tukey HSD, *Temnothorax* tandem – *Diacamma* tandem, *z* = −4.16, *P* < 0.01). The structure of *Diacamma* tandem networks was similar to the social carrying networks of *Temnothorax* due to the presence of many transportees that were carried only once (Fig. 3C). There was no significant difference in the proportion of passive workers with one incoming edge and no outgoing edge (LMM; Tukey HSD; *Temnothorax* carrying – *Diacamma* tandem, *z* = −0.32, *P* = 0.94; Fig. 3DE)

**Figure 3.**
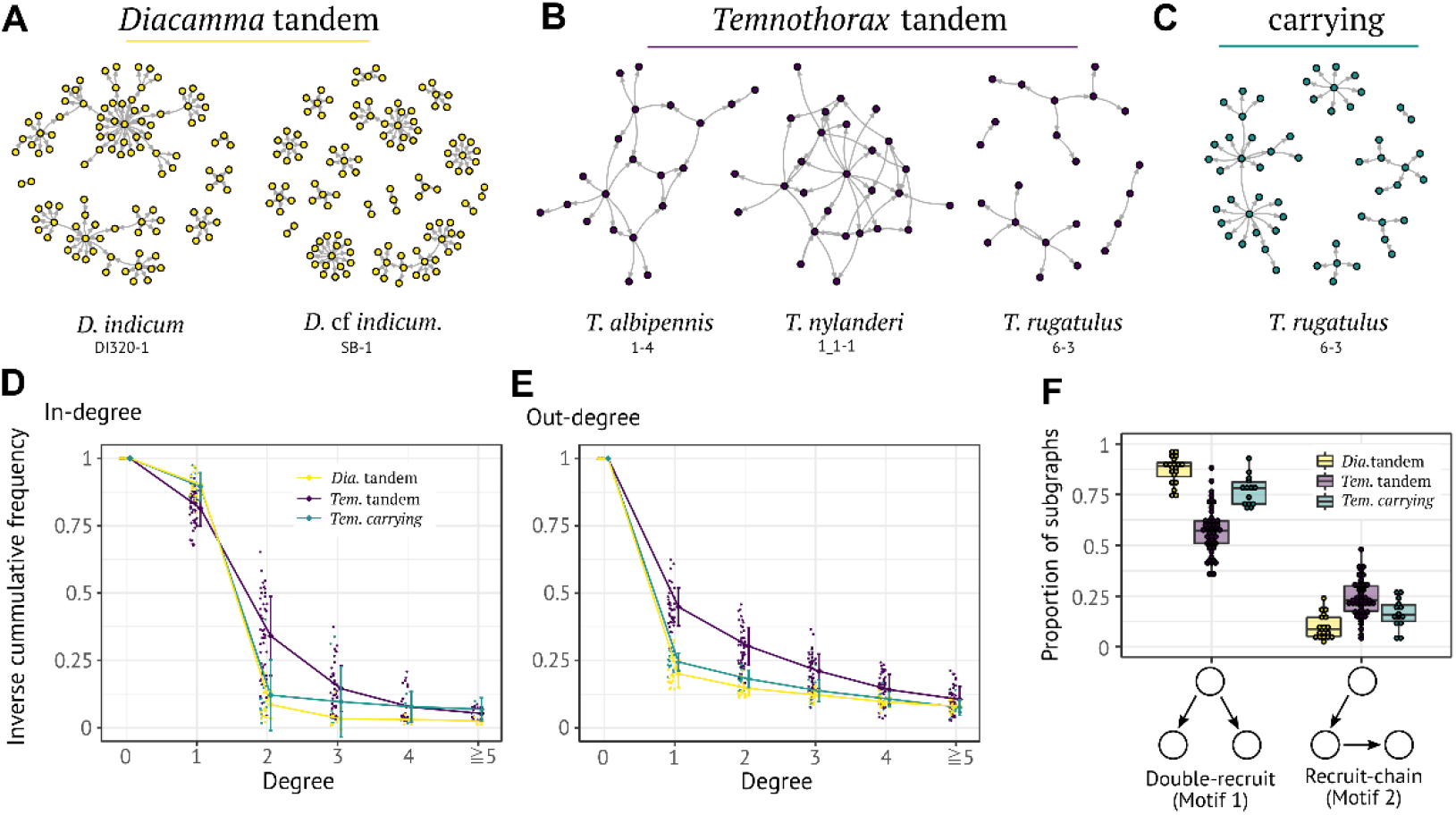
Comparison of recruitment networks in *Diacamma* and *Temnothorax* ants. (A-C) Examples of recruitment networks from different species. Nodes represent ants, and edges represent recruitments (tandem run or carrying). Colony id and emigration event are connected by a hyphen following the species name. (A) Tandem recruitment network for *D*. cf *indicum* and *D. indicum*. (B) Tandem recruitment network for *T. albipennis, T. nylanderi*, and *T. rugatulus*. (C) Carrying network for *T. rugatulus*. (D-E) Comparison of cumulative in-degree and out-degree distributions. Bars indicate mean ± s.d. (F) Comparison of the proportion of network motifs for different types of ant recruitment networks. Networks with < 20 nodes were removed from the analysis.

Network motif, a property characterizing the structure of a given network (Milo et al., 2002), provided more evidence for differences in the structure of *Diacamma* and *Temnothorax* tandem recruitment networks. We investigated the distribution of triadic motifs to identify representative patterns of information flow in recruitment networks. Among 13 possible subgraphs with three nodes, two motifs predominated, namely the “double-recruit” motif (A→B, A→C, motif 1 in (Milo et al., 2002)) and the “recruit-chain” (A→B→C, motif 2 in (Milo et al., 2002)). Double-recruit was more frequent in *Diacamma* tandem recruitment networks than in *Temnothorax* tandem networks (Tukey HSD: *Temnothorax* tandem – *Diacamma* tandem, *z* = 2.96, *P* = 0.01, Fig. 3F). On the other hand, recruit-chain was more frequent in *Temnothorax* networks than *Diacamma* networks (Tukey HSD: *Temnothorax* tandem – *Diacamma* tandem, *z* = −3.82, *P* < 0.001, Fig. 3F).

These differences are consistent with the suggestion that the tandem recruitment networks of *Temnothorax* result from individuals switching from following to leading, whereas the networks of *Diacamma* rarely contain these switches, and instead mainly represent interactions between a minority of highly active recruiters and a majority of passive followers. In contrast, we found no significant difference between motifs present in *Diacamma* tandem networks and *Temnothorax* social carrying networks (Tukey HSD: double-recruit: *z* = 0.87, *P* = 0.66, recruit-chain: *z* = −0.27, *P* = 0.96), and both represent interactions of a minority transporting the majority of other members.

### Species differences in the coordination mechanisms of tandem runs

The two members of a tandem coordinate their movement by adjusting their speed according to that of their partner. Our analysis of the changes in movement speed (i.e., the acceleration) during tandem runs showed that leaders and followers, in both *D*. cf *indicum* and *T. rugatulus*, modified their movement acceleration according to the distance to each other (Fig. S3), as previously observed in *T. albipennis* (Franks and Richardson, 2006). When the distance increased, followers accelerated to catch up with leaders, and leaders decelerated to allow the follower to reconnect. Conversely, when the distance decreased, followers decelerated, and leaders accelerated. However, there were striking differences between *D*. cf *indicum* and *T. rugatulus* in the stability and duration of tandem runs. Tandem runs of *T. rugatulus* were frequently disrupted, with all runs experiencing separation within three minutes (Fig. 4A). In contrast, tandem runs of *D*. cf *indicum* were more stable, with more than half of the runs lasting longer than 15 minutes (Fig. 4A). As a result, tandem pairs of *T. rugatulus* were separated at a distance for much longer periods than *D*. cf *indicum* over the course of observed tandem runs (Fig. 4B).

**Figure 4.**
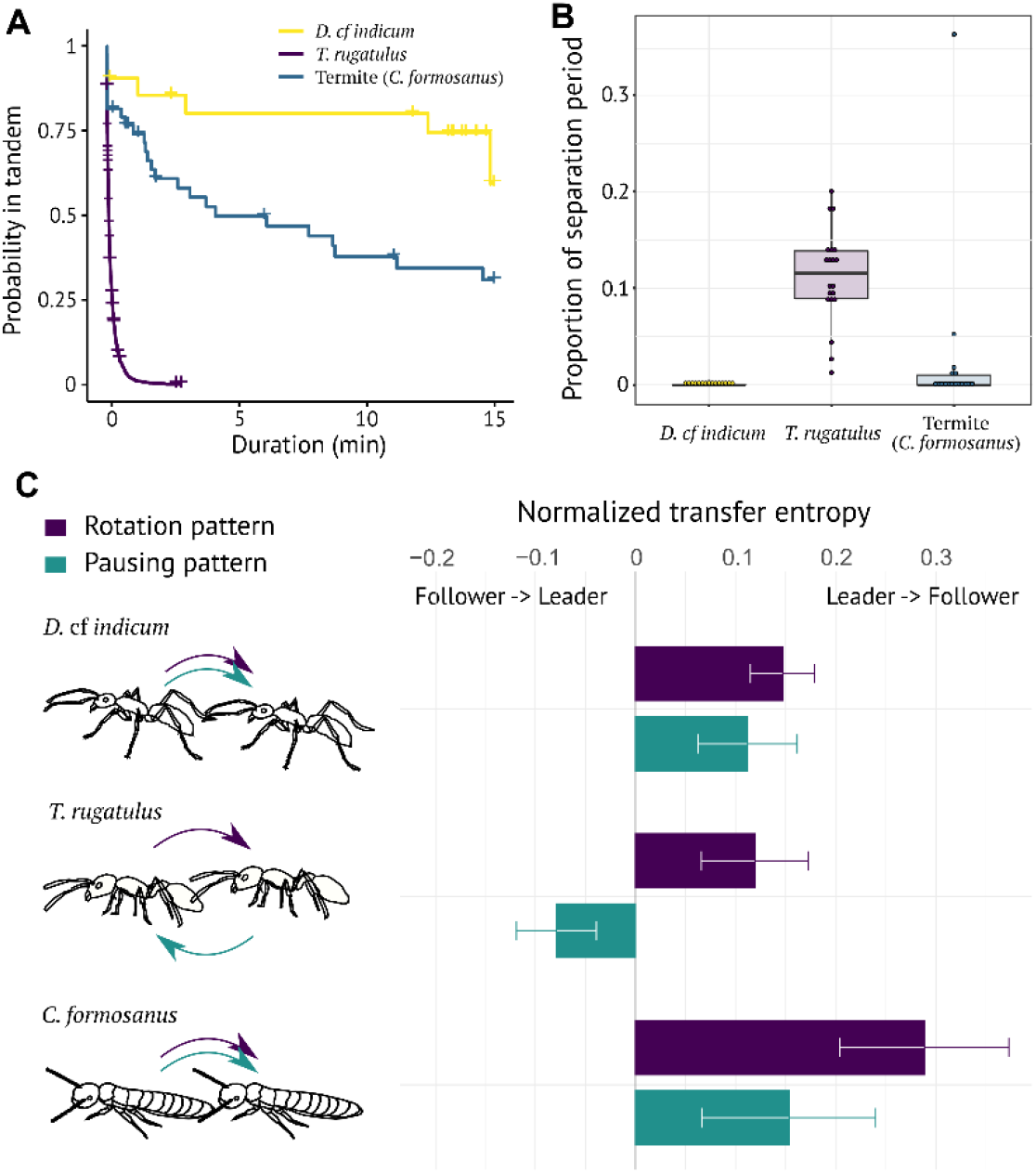
Comparison of movement coordination during tandem runs. (A) Comparison of the duration of tandem runs until interruptions. Kaplan–Meier survival curves were generated for each species. + indicates censored data due to the end of observations. There was a significant difference in duration until separation between species (Tukey HSD after mixed-effects Cox model, *P* < 0.05). (B) Comparison of the separation period during observation. (C) The predominant direction of predictive information is given by the proportion of uncertainty reduction explained by the interaction between leader and follower. Bars indicate the mean ± s.d.

The frequent interruptions of *Temnothorax* tandems are believed to play a role in information transfer. Followers trigger pauses so that they can learn visual cues that will later help them navigate the route independently. Thus, follower behavior largely drives the temporal pattern of move/pause by leaders. On the other hand, because only leaders know the route to the new nest, their behavior largely determines the movement direction of followers. This bidirectional feedback was recently formally quantified using an information theoretic analysis in *T. rugatulus* (Valentini et al., 2020b). The net flow of information between a leader and follower was quantified using transfer entropy (Lizier and Prokopenko, 2010; Porfiri, 2018; Schreiber, 2000); that is, information flow is from a leader to a follower if leader behavior predicts the future follower behavior, better than the other direction. They separately analyzed the sequence of pauses and moves, and the sequence of clockwise- and counter-clockwise turns, to examine the direction of net flow of information.

We applied the same analysis to *Diacamma* tandem runs, expecting that tandem runs in this species would lack this bi-directionality, given their greater stability and the absence of *Temnothorax*-like route-learning. Indeed, for the tandem runs of *D*. cf *indicum*, leader behavior was always a better predictor of follower behavior than the other way around, both for motion and for direction (Wilcoxon signed-rank tests, *P* < 0.01, Fig. 4C, S4). This contrasted with *T. rugatulus* tandem runs, in which leader turning behavior predicted followers turning patterns, and follower pause behavior predicted leaders’ pausing patterns (Wilcoxon signed-rank tests, *P* < 0.01, Fig. 4C). The tandem runs of *D*. cf *indicum* were qualitatively similar to those of termite mated pairs, where the present behavior of the leader significantly predicted the future behavior of the follower (Wilcoxon signed-rank tests, *P* < 0.01, Fig. 4C). This corresponds to the similar functions inferred for *Diacamma* and termites, but note that *Diacamma* tandems had fewer separation than termite tandem runs (Fig. 4AB).

## Discussion

When ants emigrate, they use recruitment communication to share information and reach a consensus on a single nest. Both *Diacamma* and *Temnothorax* ants recruit via tandem runs, a behavior which at first glance seems similar in the many ant species where it is found (Franklin, 2014). However, we demonstrated that tandem runs of these two species are functionally distinct. We found that tandem followers of *Diacamma* do not actively regulate the movement of leaders but keep consistent close contact with their leaders. This suggests that *Diacamma* prioritizes avoiding lost followers over providing opportunities for followers to collect spatial information. In *Temnothorax* ants, this mass movement of colony members is conducted by social carrying, an alternative recruitment technique, where an active ant carries a nestmate in her mandibles. *Temnothorax* instead use tandem runs to recruit additional recruiters by sharing route information to the new nest (Fig. 2). Unlike in *Diacamma, Temnothorax* followers actively regulate their leaders’ movements to facilitate route learning. This contrasts with *Diacamma* followers who usually do not need to learn the route as they remain at the goal after arrival. Thus, although both species possess a similar social behavior, there is a great deal of cryptic variation between species.

In addition to the different functions of tandem runs in *Diacamma* and *Temnothorax*, the interaction rules responsible for movement coordination are also distinct. In both genera, leaders release a short-range pheromone to help guide followers (Fujiwara-Tsujii et al., 2012; Moglich et al., 1974), and followers use tactile signals to indicate their presence to leaders. However, leaders of *Diacamma* control both the direction and speed of tandem runs, while *Temnothorax* leaders and followers consistently regulate each other’s motion (Fig. 4). In this sense, *Diacamma* tandem runs are like those used by termite mating pairs, where the female leader decides the course of movement during search for a nest site, and the male follower strives not to be separated. Note that *Diacamma* leaders modify their behavior to facilitate coordination even though the interaction is unidirectional. For example, the leader’s moving speed during tandem runs is slower than when the leader is alone (Kaur et al., 2017), contrasting with the termite leaders that show consistent movement speed even with male followers (Mizumoto et al., 2021). As a result, *Diacamma* tandem runs are highly stable with few interruptions (Fig. 4). We, therefore, suggest that *Diacamma* tandem runs should not be viewed as a simple form of recruitment by leading and following, but rather a specialized social behavior for preventing separation during emigration. Conversely, *Temnothorax* tandem runs can be seen as maximizing information sharing at the risk of frequent follower loss.

The distinct tandem runs of *Diacamma* and *Temnothorax* presumably reflect different selective pressures posed by their nest-moving contexts. Ant colonies face a speed-accuracy trade-off during nest site selection (Nigel R Franks et al., 2003), where the nest quality should be important in *Temnothorax*, while the highest priority should be speed for *Diacamma* emigration. In *Temnothorax* ants, colonies live in preformed cavities such as hollow nuts or rock crevices, rather than building their own nests. This places a premium on their ability to find and select a high-quality nest from the available options (Dornhaus et al., 2004). Site quality may be critical for colony defense from competitors (Nigel R. Franks et al., 2003), and suitable nest sites are limited resources (Herbers, 1986) for which colonies compete (Foitzik and Heinze, 1998). Therefore, *Temnothorax* workers rely on tandem runs for information sharing and collective decision-making (Sasaki et al., 2015). In contrast, *Diacamma* ants live in rainforests, where heavy rains result in nest flooding and demand rapid evacuation to temporary sites (Kolay and Annagiri, 2015a). Colonies frequently move (Fukumoto and Abe, 1983), at an estimated interval of every 1-2 weeks (Win et al., 2018). Stable and fast tandem runs would be favored because they facilitate prompt and flexible colony movement, where workers rely much more on individual decisions than collective decision-making (Anoop and Sumana, 2015). The tandem runs of *Diacamma* are made even more efficient by the followers’ habit of carrying brood or food items (Kaur et al., 2017) (Fig. 1B); this has rarely been observed in *Temnothorax*, possibly due to its potential interference with route-learning. Furthermore, although emigrating *Diacamma* colonies prefer better nest sites (Sahu et al., 2019), they can excavate their own underground nests from pre-existing openings on the ground (Bhattacharyya et al., 2021; Kaur et al., 2012), and thus are less limited by nest site availability than *Temnothorax*.

Lineage-specific selective pressures may also be at work in other independent origins of tandem running behavior besides *Temnothorax* and *Diacamma*. For example, recruitment speed may also be important for *Pachycondyla harpax* as they need to rapidly exploit or defend food sources in their highly competitive tropical habitat (Glaser et al., 2021). Route-learning may be especially important for *Harpagoxenus sublaevis* ants that use tandem runs to recruit nestmates for slave raids (Buschinger et al., 1980). In this species, multiple tandem followers go on to lead other ants to the target nest, which requires them to learn the route. *Camponotus*, on the other hand, may differ from both *Diacamma* and *Temnothorax*. For example, tandem followers of *C. consobrinus* already have navigational knowledge of the destination but use tandem runs to potentially improve their spatial information about the complex branching structure of their arboreal foraging grounds (Schultheiss et al., 2015). This is distinct from the route-learning of *Temnothorax* ants, whose followers are typically naïve. Furthermore, interspecific differences within a genus are also possible (Alleman et al., 2019; Möglich, 1978; Pratt, 2005). The cryptic diversity in pair movement coordination can be understood through comparative behavioral analysis, aided by our semi-automatic system to extract tandem run coordination from a group of ant foragers (Methods, Fig. S6).

In *Diacamma* tandems, most followers do not become leaders, but a few followers turn into leaders after tandem runs (Fig. 2BC and 3AF). Also, in natural conditions, leaders also follow tandem runs to ensure the best location of new nest sites when there are multiple options (Kaur et al., 2012). Thus, followers of *Diacamma* ants might acquire some information through tandem runs. How do *Diacamma* leaders know the location of the new nest if their tandem runs lack a *Temnothorax*-like route-learning function? There are two non-exclusive explanations. First, tandem followers of *Diacamma* might be able to learn the route differently from *Temnothorax* followers. Even without frequent interruptions, as observed in *Cataglyphis* desert ants, followers could collect spatial information through a stride integrator that accounts for stride number and the respective stride length (Wittlinger et al., 2006) or optic flow of image motion experienced during travel (Pfeffer and Wittlinger, 2016). Second, *Diacamma* leaders might have knowledge of landscapes surrounding the nests before the colony emigration begins, allowing them to become leaders without step-by-step route-learning as in *Temnothorax* ants. For example, honeybee foragers explore their nest surroundings before searching for food sources to establish familiarity with the landmarks making their home(Menzel et al., 2000). In *Diacamma*, leaders may already know the locations of candidate nest sites because tandems are primarily led by foragers (Fukumoto and Abe, 1983), who increase exploring activity during rainy seasons (Win et al., 2018). Note that a similar mechanism can also be partly used in *Temnothorax* ants (Stroeymeyt et al., 2011, 2010). The functional capacity of ant tandem running behavior strongly depends on the cognitive ability of ant species, which could be a source of the diversity of this social behavior.

## Conclusion

Convergent evolution is often considered an outcome of similar evolutionary pressures occurring in different lineages. However, similarity in phenotype does not always guarantee the same functional capacities in natural history (Losos, 2011). Similar morphology has been associated with different functions among different lineages, as evidenced in the tree growth habit (tall plants with a thickened single trunk (Donoghue, 2005)), foot webbing in salamanders (Jaekel and Wake, 2007), and saber-shaped teeth in carnivores (Lautenschlager et al., 2020). We show that this is even true for animal social behavior. Tandem running behaviors of *Temnothorax* ants and *Diacamma* ants play different functional roles during colony emigration, and similar movement coordination comes from different forms of interactions. By quantifying the communication behavior of social animals in a comparative framework, this study will improve our ability to resolve difference in the evolution of social behavior in a wide variety of lineages.

## Materials and Methods

### Ant recruitment dataset

To examine recruitment dynamics in ant tandem runs, we used datasets for leader-follower combinations of tandem runs during colony emigration. In *Diacamma*, recruitment is only via tandem run, while *Temnothorax* recruitment can be via tandem or social carrying. Only tandem recruitment data were available for *D*. cf *indicum* from Japan, *D. indicum, T. albipennis*, and *T. nylanderi*. Both tandem run and social carrying data were available for *T. rugatulus*. Note that we treated *D. indicum* and *D*. cf *indicum* separately because of their locational segregation and historical distinction (Tsuji, 2021). *D. indicum* is reported from the eastern and southern parts of India and Sri Lanka (Viginier et al., 2004). *D*. cf *indicum* refers to an ant population, closely related to *D. indicum* but separated in a distance. This population is from Okinawa Island, Japan, has long been reported as “*Diacamma* sp. from Japan,” and is still awaiting a formal taxonomic description (Shimoji and Dobata, 2022). The context of each emigration experiment is as follows:

#### (i) *Diacamma* cf *indicum*

We used four colonies of *D*. cf *indicum* collected in Onna and Naha, Okinawa, Japan, between August and October 2021 (colony ID: K4, K8, SA, SB; colony size: 115, 127, 111, 129 workers + one gamergate, respectively). Each colony was maintained in an experimental nest, a plastic container (24.1 cm × 14.1 cm × 3.7 cm) fill with plaster and covered with a translucent red plastic plate. Until the experiment, the nest was connected to a foraging area (41 cm × 26 cm × 7 cm) that was provided with a water tube and food (mealworms and standard artificial diets (Dussutour and Simpson, 2008)) under laboratory conditions (25 ± 1 °C with stable light-dark cycle 14: 10 light: dark). We marked all workers by using a unique combination of four colors for individual identification.

Colony emigration was observed in a square arena (100 cm × 100 cm × 30 cm) filled with sand collected from the field (Kwansei Gakuin University, Japan). The old and new nests were positioned at 55 cm. To inform the colony about the environment of the experimental arena, we kept the colony in this arena for three days with sufficient food and water. The new nest was unroofed during this period and thus unavailable for the colony. After three days, we moistened and covered the new nest to make it available. As we did not add water to the old nest while maintenance, the old nest was drier than the newly moistened nest, whereas the new and wet nest was preferable for *Diacamma* colonies. We also removed the red plate on the old nest to induce colony emigration. A digital video camera (HC-V480MS, Panasonic, Japan) was located at the top of the entrance to the new nest, and we recorded the time of each tandem run during the nest relocation and the IDs of each leader and follower.

#### (ii) Diacamma indicum

Experiments with *D. indicum* were performed as part of a study on tandem leader removal during multiple emigrations (Kolay and Annagiri, 2015b). Sixteen colonies of *D. indicum*, collected from Mohanpur, West Bengal, India, between September 2013 and May 2014 (colony ID: see Fig. S8; colony size: 133.9 ± 37 adults [average±s.d.] + one gamergate), were used in the experiments. All workers had unique colored paint marks. The colony emigrated from the old and broken nest to the identical new nest. Colony emigrations were observed over a wooden bridge (1520 mm) that connected the old and new nests. Each colony emigrated twice: an initial control emigration and the following leader or random member removal experiment. We used data from the control emigration for this study. We recorded the time and the participating workers of all tandem runs during colony emigration. Further details can be found in (Kolay and Annagiri, 2015b).

#### (iii) Temnothorax rugatulus

Experiments with *T. rugatulus* were performed as part of a study on the division of labor during multiple emigrations (Valentini et al., 2020a). Three colonies of *T. rugatulus*, collected in the Pinal Mountains near Globe, Arizona, in 2009 (colony ID: 6, 208, 3004; colony size: 78, 81, and 33 workers + one queen, respectively), were used in the experiments. All workers had unique colored paint marks. The colony emigrated from an old and broken nest to either of two new candidate nests with different quality (good and mediocre). The two new nests were 50 cm from the old nest in a rectangular arena (37 cm× 65 cm). Each colony emigrated five times with a rest interval of two to five days between emigrations. The time and involved workers were recorded for all tandem running and social carrying events during colony emigration. Further details can be found in (Valentini et al., 2020a).

#### (iv) Temnothorax albipennis

Tandem running interaction in *T. albipennis* was recorded in experiments to study collective decision-making (Richardson et al., 2018). Six colonies of *T. albipennis*, collected on the Dorset coast, UK, during 2011 (colony ID: 1-6; colony size: 72-113), were used in the experiments. All workers were tagged with RFID micro transponders (PharmaSeq, NJ, USA) for individual identification. The colony emigrated from a nest of poor quality to either of two better nests of identical quality. These three nests were placed in a rectangular arena (45-75 cm) with an equilateral-triangle arrangement. The emigration procedure was repeated five times for each colony at seven-day intervals. The time and involved workers were recorded for all tandem running events during colony emigration. Further details can be found in (Richardson et al., 2018).

#### (v) Temnothorax nylanderi

Tandem running interactions in *T. nylanderi* were recorded in leader or follower removal experiments (Richardson et al., 2021). Twelve colonies of *T. nylanderi*, collected in the Forêt de Dorigny, Switzerland, between June and October 2018, were used in the experiments (colony ID: 1_1-3_4; colony size 71-116 workers). All workers had unique colored paint marks. The colony emigrated from a low-quality and half-broken nest to either of two new nests with better and identical quality. These three nests were placed in a rectangular arena (46-78 cm) with a triangle arrangement. Each colony was subjected to five emigrations at one-week intervals between successive emigrations. Specialized leaders or followers were removed before the fifth emigration. Thus, we used the data for the 1^st^ through 4^th^ emigrations in this study. Participating workers were recorded for all tandem runs during colony emigration. Further details can be found in (Richardson et al., 2021).

### Comparison of recruitment dynamics

Information on the whole emigration was available in *D*. cf *indicum, D. indicum*, and *T. rugatulus*, while only tandem run (initial stage) information was available in *T. albipennis* and *T. nylanderi*. We first compared the proportion of individuals engaged in each recruitment activity between *D*. cf *indicum, D. indicum*, and *T. rugatulus*. We used generalized linear mixed models (GLMMs) with binomial errors and a logit link function. The recruitment type was treated as an explanatory variable, and the emigration event was included as a random effect (random intercept). The likelihood ratio test was used to assess the statistical significance of the inclusion of each explanatory variable (type II test). The GLMM was used for each of six pair-wise comparisons of four different recruitment types. The analysis was performed for the proportion of leaders and followers separately. GLMM analyses were conducted using the ‘car’ and ‘lme4’ packages (Bates et al., 2015).

We next investigated the behavioral sequence of individuals during colony emigrations for each species. The behavioral states of *Diacamma* (or *T. albipennis* and *T. nylanderi*) included three states: tandem leader, follower, and end of the colony emigration, while *T. rugatulus* included five states: tandem leader, follower, social carrier, being carried, and end of the colony emigration. We calculated the probability of transitioning from one state to another for each state. Passive individuals (tandem followers or being carried) can become active recruiters (tandem leaders or social carriers) after recruitment. Using Fisher’s exact test, we compared the probability of a passive ant being activated across species.

### Recruitment network analysis

For each emigration event and recruitment tactic, we obtained 125 networks (*D*. cf *indicum*: 4, *D. indicum*: 16, *T. albipennis*: 30, *T. nylanderi*: 48, *T. rugatulus* [tandem]: 13, *T. rugatulus* [social carrying]: 14). The sequence of tandem runs (or social carrying) was represented as a static network. Ants who participated in recruitment were represented as nodes, and tandem runs were represented as directed links pointing from the leader (or carrier) to the follower (or being carried). Note that no tandem was observed in one emigration event in *T. rugatulus*. Networks with < 20 nodes were removed from the analysis.

First, we examined the in-degree and out-degree distributions to compare the overall network structures between *Diacamma* and *Temnothorax* tandem recruitment structures. We compared these distributions using Kolmogorov-Smirnov (KS) test (data were pooled for each genus). Also, we compared the proportion of workers with one incoming edge and no outgoing edge between recruitment types. We used the LMM similar to above, after logit transformation of the proportion data by adding 0.01 to the observed proportions to avoid infinite values (Warton and Hui, 2011). Note that these parameters may not always follow normal distributions, but LMM is robust against violations of distribution assumptions (Schielzeth et al., 2020).

Next, we investigated the motifs of these structures to compare the property of network structures. Network motifs are overrepresented small subgraphs in a given network (Milo et al., 2002). The motif property is reported to be independent of colony size in other social insect research (e.g., (Waters and Fewell, 2012)) and thus ideal for our comparison between species. There are 13 possible directed subgraphs for three-node patterns. We used the function graph.motifs() in the R package ‘igraph’ (Csardi and Nepusz, 2006) to obtain the number of each subgraph for each network. Then, we compared the proportion of two specific patterns (motifs 1 and 2), which were overrepresented in recruitment networks, between lineages and recruitment types (*Diacamma* tandem, *Temnothorax* tandem, and *Temnothorax* carrying).

The proportion of subgraphs was compared between lineages and recruitment types (*Diacamma* tandem, *Temnothorax* tandem, and *Temnothorax* carrying). We used linear mixed models (LMM), where the recruitment type was treated as an explanatory variable, and the colony nested within species was included as a random effect (random intercept). In cases of significant effects, we ran Tukey’s post hoc test using the glht() function in ‘multcomp’ package (Hothorn et al., 2008). Note that the size of the network (= number of nodes) was different between recruitment types, and we confirmed that there was no relationship between the number of nodes and the proportion of each motif (Fig. S7).

### Tandem trajectory dataset

We used movement trajectories of leaders and followers in tandem pairs from two ant species, *D*. cf *indicum* and *T. rugatulus*, and two termite species, *Coptotermes formosanus* and *Reticulitermes speratus*. The tandem observation experiment for each species is as follows:

#### (i) *Diacamma* cf *indicum*

We used four colonies of *D*. cf *indicum* (colony size:136-160) collected in Onna and Nakijin, Okinawa, Japan, in August 2021. Each colony was maintained in a humid artificial nest covered by a red plastic plate (14.8 cm × 8.4 cm × 3.2 cm) connected to a foraging area (24 cm × 18 cm × 8.5 cm). To obtain sufficiently long trajectories of tandem runs, we prepared a large experimental arena (100 cm × 100 cm × 30 cm) with a twisting bridge connecting the old nest to a new nest. The bridge consisted of 12 straight paths in parallel with 10 U-turns connecting two straight paths (Fig. S5). The design and dimensions of this arena were informed by a previous study on *T. rugatulus* (Valentini et al., 2020b). A digital video camera (HC-VZX992M, Panasonic, Japan) was located at the top of the arena, and we recorded the whole emigration at 30 frames per second with FHD resolution. For each colony, we selected 10 (or 16 for one colony) tandem runs that were longer than 5 minutes. We subdivided the whole video into multiple clips so that each clip includes one selected tandem running event. In total, we obtained 46 video clips.

Extracting trajectories of a leader and a follower in a tandem running pair is challenging because partners are often in contact with each other. Most tracking software recognizes a tandem pair as a single individual and fails to distinguish partners from each other. The coordinates of tandem partners can be well extracted using UMATracker software (Yamanaka and Takeuchi, 2018), as evidenced by previous studies (Mizumoto et al., 2022, 2021, 2020; Mizumoto and Dobata, 2019; Valentini et al., 2020b). One of the prerequisites of the UMATracker is a fixed number of individuals in a video frame. However, this is not the case in ant colony emigration experiments, where ants continuously appear and disappear in the arena by entering or coming out of the nests (Fig. S6A). A previous study manually subdivided the video and trimmed the frames so that one video clip included a fixed number of ant individuals (Valentini et al., 2020b). As this process is time-consuming, we needed an alternative solution to efficiently obtain data from multiple species. To overcome this challenge, we established a system to track ant tandem running behavior semi-automatically by combining another tracking software, FastTrack (Gallois, 2021), with UMATracker (Fig. S6). Although FastTrack cannot recognize a tandem running pair as two different partners, it can handle a changing number of objects in a region of interest, which is suitable for analyzing ant colony emigration events.

Using FastTrack, we first recognized a tandem running pair as a single object and extracted the rough trajectory of the centroid of the pair (Fig. S6B). Because FastTrack skips any frame in which it failed to recognize an object, we gave tentative coordinates of the tandem in skipped frames by using R to calculate the averaged values of the frames before and after the skipped frames. Next, from the tentative coordinates of tandem runs, we created a video that showed only the focal tandem running pair, using the OpenCV library and Python scripts. We did this by making a background frame by averaging every 1,000 frames throughout the clips. Then, we covered the video clips with this background frame except for a 20 × 20 pixel area surrounding the focal tandem coordinates; this masked all individuals other than the focal tandem pair (Fig. S6C). Finally, we used UMATracker on this masked video clip to extract the trajectories of the tandem leader and follower (Fig. S6D). This process inevitably missed pairs in the masked video clip when they were separated by more than ∼20 pixels. To analyze these, we applied UMATracker to the original videos for these frames by manually specifying the region of interest. Finally, our videos sometimes missed tandem running pairs that went behind the bridge (46 events, 25,209 frames in total, ≈ 2.20% of all frames) or due to the blackout between two successive video clips imposed by our video camera (13 events, 668 frames in total, ≈ 0.06% of all frames). For these exceptional frames, we estimated the coordinates by assuming that ants were moving linearly and at a constant speed. The Python script for this process is available in the datasets.

As a result, we obtained 46 trajectories of the centroid of the leaders and followers at 30 frames per second (1,146,126 frames). Among them, 15 are longer than 15 minutes, and 29 are longer than 10 minutes. Trajectories were converted from pixels to millimeters using a scaling factor estimated by measuring known features of the experimental arena with ImageJ (Schneider et al., 2017). The body length was also measured using ImageJ (mean ± s.d. = 10.54 ± 0.86 mm).

#### (ii) Temnothorax rugatulus

Trajectories of tandem leaders and followers were available from previously published work on *T. rugatulus* (Valentini et al., 2020b). Six colonies (colony size: 30-60) were induced to move from an old nest to a new nest located at a diagonal position in a large experimental arena (370 × 655 mm). The arena was subdivided by five barriers (10 mm by 310 mm), forming a twisting path to observe sufficiently long tandem runs. The colony emigration was recorded once for each colony at 30 frames per second using a video camera with 1K resolution. One to six tandem runs that lasted more than 15 min were selected for each colony, and the trajectories of the leader and the follower were extracted using the UMATracker software platform. The centroids of each runner’s body were tracked at 30 frames per second. Trajectories were converted from pixels to millimeters using a scaling factor estimated by measuring known features of the experimental arena with ImageJ (Schneider et al., 2017). The body length was also measured using ImageJ (mean ± sd = 2.34 ± 0.3 mm).

#### (iii) Termites

Trajectories of tandem leaders and followers were available for *Reticulitermes speratus* and *Coptotermes formosanus* (Mizumoto and Dobata, 2019). Two colonies with alates of *C. formosanus* were collected in Wakayama, Japan, in June 2017, and five colonies of *R. speratus* were collected in Kyoto, Japan, in May 2017. After a controlled flight in the lab, dealates (individuals that had shed their wings) were used for tandem run experiments. A female and a male termite were introduced in the experimental arena (a Petri dish filled with moistened plaster), and their behavior was recorded for 60 minutes. In total, 17 pairs were observed for *C. formosanus* and 20 pairs for *R. speratus*. The resolution of the videos was reduced to 640 by 480 pixels for movement tracking. The centroids of each runner’s body were tracked at 30 frames per second. Trajectories were converted from pixels to millimeters by using as a scale the diameter of the arena petri dish (145 mm). We used the body length measured in (Valentini et al., 2020b) (*C. formosanus*: mean ± sd = 8.89 ± 0.42 mm, *R. speratus*: mean ± sd = 5.5 ± 0.3 mm). In this study, we only present the results of *C. formosanus* in the main figures because they showed a qualitatively similar pattern compared to ant tandem runs.

### Trajectory analysis

We down-sampled all videos to a rate of five frames per second (FPS) (= every 0.2002 s) for the analysis in this section. To compare the stability of tandem runs between species, we defined two individuals to be in tandem when the distance between their centroids was less than two body lengths (Valentini et al., 2020b). As a result, we obtained 21, 1523, 43, and 89 tandem run events for *D*. cf *indicum, T. rugatulus, C. formosanus*, and *R. speratus*, respectively. We compared the duration of tandem runs using the mixed-effects Cox model (coxme() function in the ‘coxme’ package in R (Therneau, 2015)), with species as a fixed effect and pair id as a random effect. The random effect accounted for the inclusion of multiple tandem events for each pair of termites. The likelihood ratio test was used to determine the statistical significance of each explanatory variable (type II test). Observations interrupted by the end of the video were treated as censored data.

### Information transfer between tandem leaders and followers

The parallel information flows within the same behavior can be obtained by extending transfer entropy to different symbolic representations of the same trajectory datasets (i.e., patterns embedded in symbols representing the direction of motion or the speed of motion) (Valentini et al., 2020b). We applied the same methodology to the tandem running behavior of *D*. cf *indicum* to compare it with previously published analyses of *T. rugatulus* and termites (refer to (Valentini et al., 2020b) for a detailed description of this methodology).

We first discretized trajectories of each runner to obtain time series describing the pausing and rotation patterns (Valentini et al., 2020b). For the pausing pattern, the behavior of each runner was either of two states: pause (P) or motion (M). For the rotation pattern, it was either clockwise (CW) or counter-clockwise rotation (CCW). The P/M states were distinguished from each other using the threshold for step length, the distance traveled by a runner between two successive frames. The threshold was set as the 10th percentile of the probability distribution of step length (Valentini et al., 2020b). A step length shorter than the threshold was represented as P; otherwise was represented as M. The threshold was obtained separately for each species and sampling period. The CW/CCW states were distinguished based on the direction of motion computed as the cross-product of movement vectors between successive frames. If no rotation was detected (i.e., cross-product equal to 0), the rotation direction was copied from the previous time step.

Given that *L* and *F* are behavioral sequences of the leader and the follower, transfer entropy from *L* to *F* is defined as

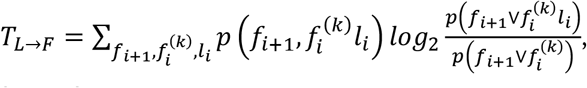

where *l*_*I*_ and *f*_*i*_ are the values of sequence *L* and *F* at time *i*, respectively, and 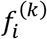 is the *k*-history of *F* at time *i* (i.e., the last *k* states in the sequence). We computed the transfer entropy in both directions, *T*_*L*→*F*_ and *T*_*F*→*L*_, and compared them to identify the predominant direction of information flow. The difference in transfer entropy between the two directions, *T*_*L*→*F*_ − *T*_*F*→*L*_, is called net transfer entropy (Porfiri, 2018; Valentini et al., 2020b). The value is positive when information flow from leader to follower is predominant (*T*|*L* → *F* > *T*_*F*→*L*_) and negative when flow from follower to leader (*T*|*L* → *F* < *T*_*F*→*L*_) predominates.

For comparison purposes, we processed transfer entropy as follows. First, to ensure that the results were not an artifact of finite sample size, we artificially created surrogate datasets by randomly pairing time series of leaders and followers from different tandem pairs (Porfiri, 2018; Valentini et al., 2020b). We computed transfer entropies for these datasets and discounted their mean from experimental data for data correction. This process enables us to focus on the causal interaction between tandem partners because leaders and followers from different pairs lack causal interactions by definition but are influenced by the same environmental cues of the experimental arena. We generated 50 surrogate datasets for each species and parameter configuration. Additionally, we obtained normalized transfer entropy by dividing it by its maximum value (Porfiri, 2018; Valentini et al., 2020b). Normalized transfer entropy ranges from 0, when the leader and follower are independent of each other, to 1, when the follower behavior is entirely determined by the leader behavior.

There are two parameters in our information-theoretic analysis, the sampling period of continuous spatial trajectories and the history length of transfer entropy, *k*. The parameter combinations that maximize the efficiency of detecting information flow can be variable across species or behaviors (Valentini et al., 2020b). To find optimal parameters, we computed net transfer entropy for 900 different parameter configurations for each species (history length *k* ∈ {1,…, 20} and sampling period {0.0334s, …, 1.5015s}). The resulting landscapes of net transfer entropy show robustness to different parameter values over most of the tested range (Fig. S4). We selected the parameter configurations that maximize the net transfer of information. For the chosen parameter configurations, we performed two statistical tests. First, we tested if the experimental data showed significantly greater values of transfer entropy with respect to the surrogate data. We used one-sided two-sample Wilcoxon rank-sum tests with continuity correction. Second, we tested differences in the flows of information between the two possible directions (from leaders to followers and from followers to leaders) to determine which among the leader and the follower was the predominant source of information. We used one-sided paired Wilcoxon signed-rank tests with continuity correction. All information-theoretic measures were computed using the ‘rinform-1.0.1’ package in R (Moore et al., 2018).

All of the data analysis was performed using R v4.0.1. (R Core Team, 2020). The R scripts and data are available in the datasets.

## Authors contributions

N.M. Conceptualization, methodology, software, validation, formal analysis, data curation, writing-original draft, writing-review & editing, visualization, supervision, project administration, funding acquisition

Y.T. Conceptualization, methodology, formal analysis, investigation, resources, data curation, writing-review & editing

G.V. Conceptualization, methodology, software, resources, writing-review & editing

T.O.R. Resources, writing-review & editing

S.A. Resources, writing-review & editing

S.C.P. Conceptualization, methodology, writing-review & editing, supervision

H.S. Conceptualization, methodology, validation, formal analysis, investigation, resources, data curation, writing-review & editing, supervision, project administration, funding acquisition

## Acknowledgments

We are grateful to Yusuke Ishizuka, Akari Nishina, Riio Yamashita, and Pan Zilin for their assistance with field collection. This study is supported by Grant-in-Aids for Scientific Research (B) to S.H. and N.M. (grant number: 22H02364), for Transformative Research Areas (B) to S.H. (22H05068), for JSPS Fellows to N.M. (20J00660), and for Early-Career Scientists to N.M. (21K15168). N.M. was supported by a JSPS Research Fellowship for Young Scientists CPD, supervised by T. Bourguignon.

## Supplementary Text

### Phylogenetic comparative analysis

We summarized the information about ant tandem running behavior at the genus level. If at least one species in the genus shows tandem running behavior in any context (recruitment to food, nest, or slave-hunting sites), we counted that genus has tandem running behavior. Almost all of the information comes from (Reeves and Moreau, 2019), which compiles the information about foraging modes in ant species. As (Reeves and Moreau, 2019) did not consider tandem running behavior during colony emigration, we newly included *Diacamma* (used in this study) and *Hypoponera* (Franklin, 2014) as genus that performs tandem running. We considered that tandem running is absent in the genus with information on foraging modes (Reeves and Moreau, 2019) but without tandem running information.

We modified the species-level ant phylogeny reconstructed by (Matthew P Nelsen et al., 2018). This phylogenetic tree included 1730 ant species from all extant subfamilies and 317 of the 334 extant genera. Because species in this phylogeny did not entirely match with the species with tandem or foraging information, we reduced the tips to build a genus-level phylogenetic tree. Note that this phylogeny randomly selected one representative species for every genus, and thus the polyphyletic genera, such as *Tetramorium*, were represented as monophyletic groups occupying singular phylogenetic positions. We also removed genera without information on foraging mode (Reeves and Moreau, 2019). As a result, the tree included 161 genera (Fig S1).

We estimated marginal ancestral states of tandem running behavioral ability in ants, using the function rerootingMethod() in R package ‘phytools’ (Revell, 2012). We used a maximum likelihood model with an equal rate of transition among states.

**Figure S1.**
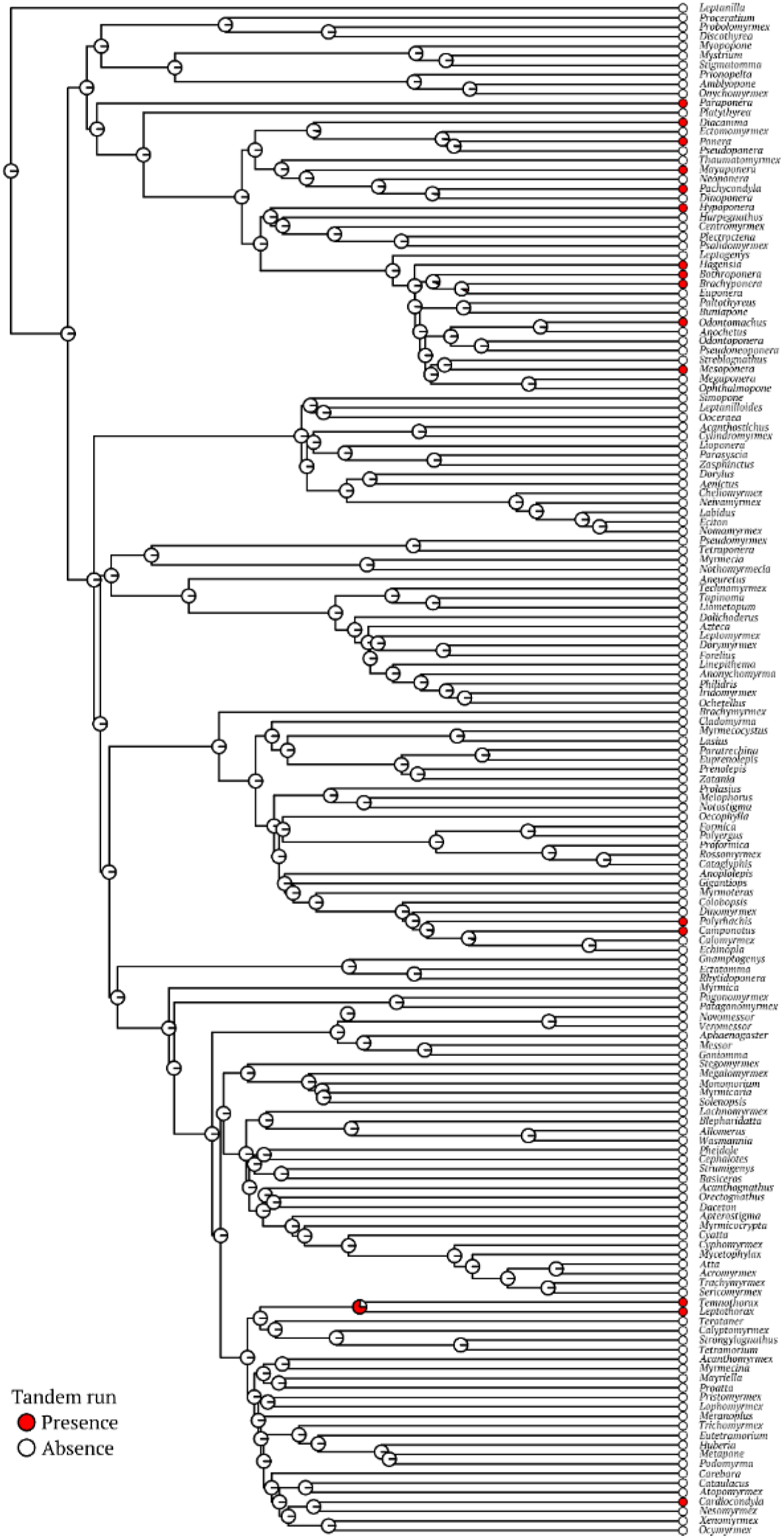
Genus-level ant phylogeny along with tandem information.

**Figure S2.**
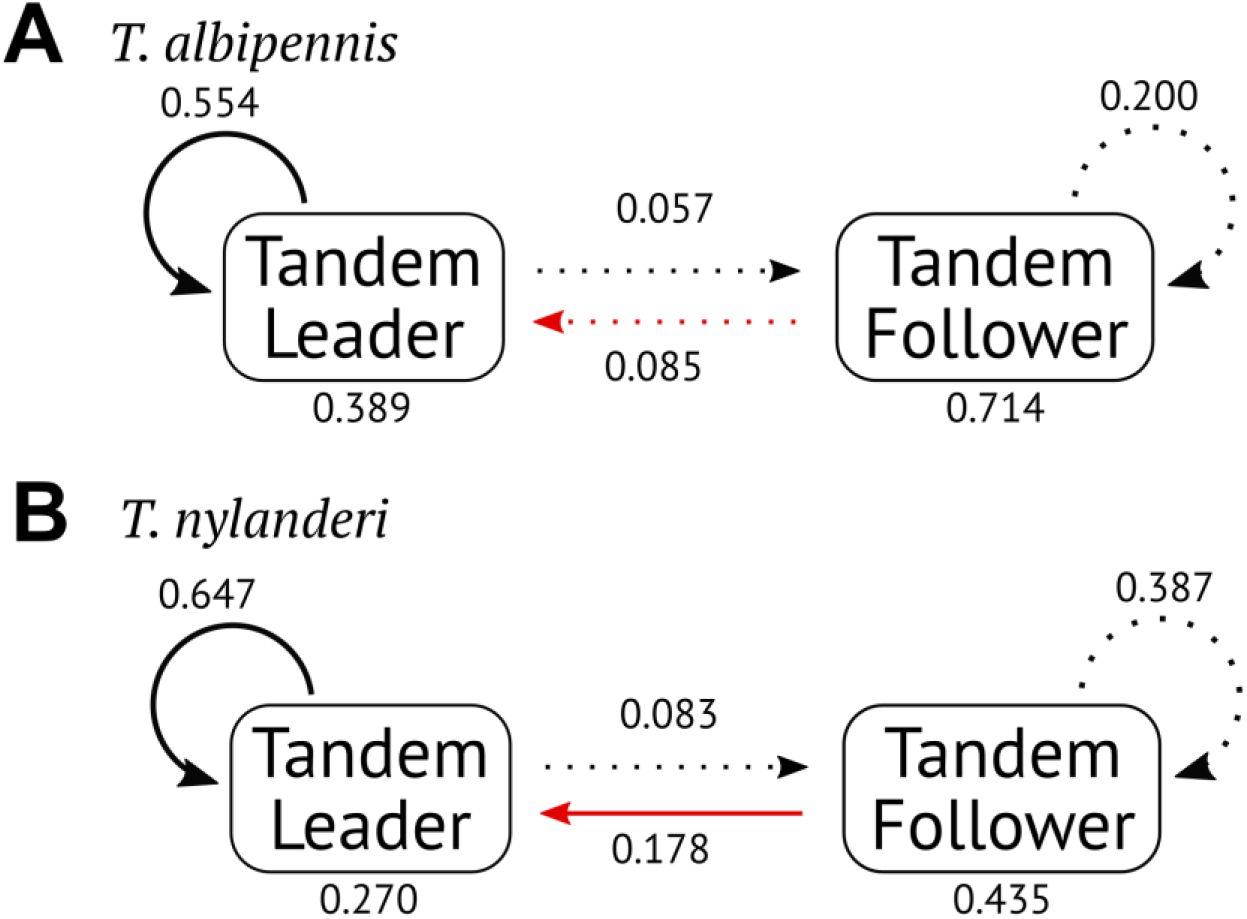
Transition diagram of worker tasks during a nest emigration event in (A) *T. albipennis* and (B) *T. nylanderi*. Recursive arrows indicate performing the same task in a row, while the numbers below the boxes indicate the probability of concluding the colony emigration after the task. Transitions with probability < 0.2 are in dashed lines, while those with probability < 0.05 are not shown in the figure. Red lines indicate transition from a passive role to an active role, suggesting successful recruitment. The probability of being recruited in *T. albipennis* tandem was significantly lower than *T. rugatulus* and *T. nylanderi* (Fisher’s exact test, *P* < 0.01), but not different from *Diacamma* tandems or *T. rugatulus* carrying (*P* > 0.2). That of *T. nylanderi* tandem was significantly different from any others (P < 0.01), which was lower than *T. rugatulus* tandem, but higher than others.

**Figure S3.**
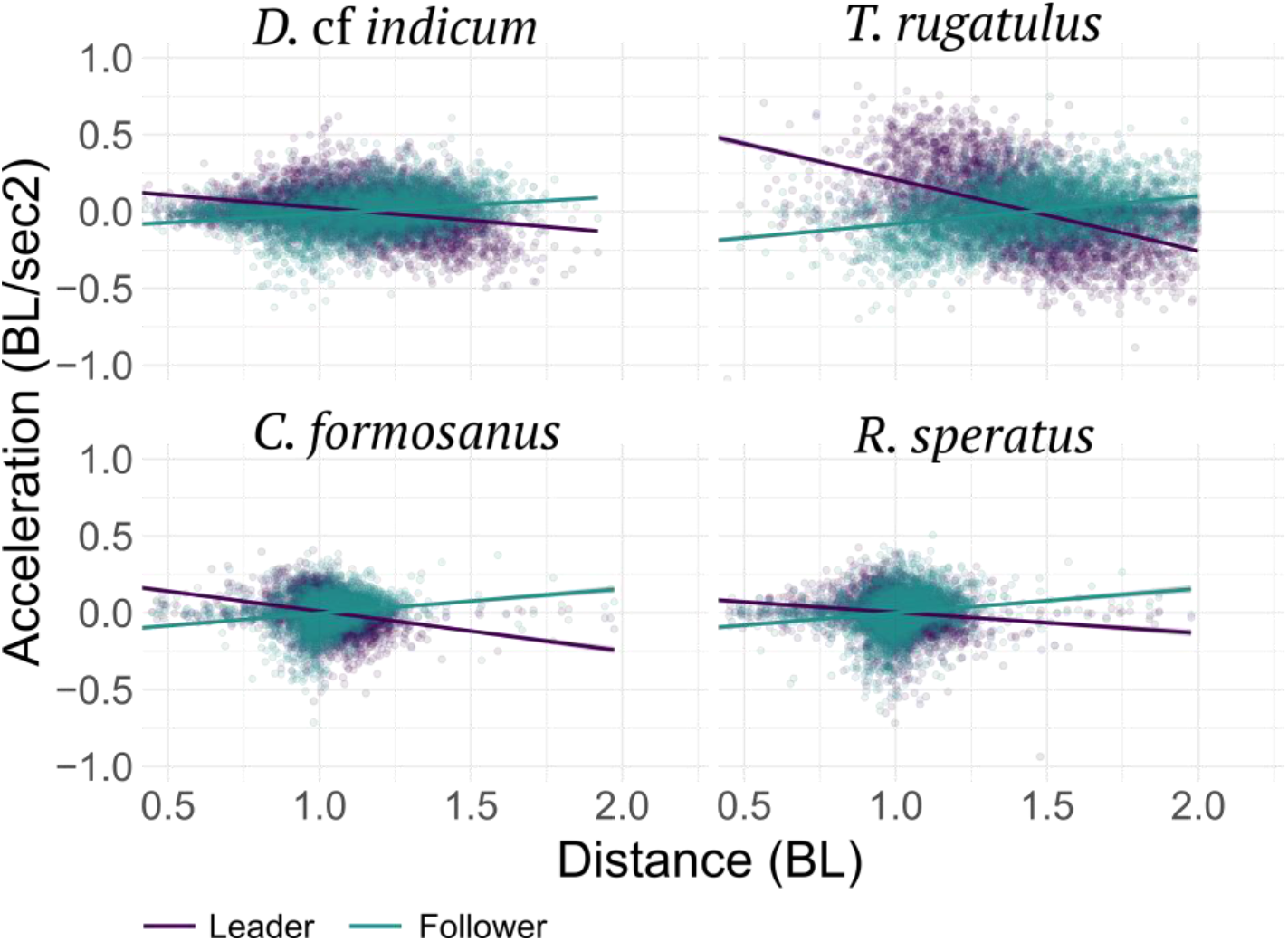
Regulation of motion acceleration during tandem runs. Plots were reduced to 5,000 data points randomly selected from each whole dataset. Slopes were significantly greater than 0 in followers (LMM with tandem pair as a random effect, *P* < 0.001, *D*. cf *indicum*: estimate = 0.126, *T. rugatulus*: estimate = 0.197, *C. formosanus*: estimate = 0.204, *R. speratus*: estimate = 0.164), while slopes were smaller than 0 in leaders (LMM with tandem pair as a random effect, *P* < 0.001, *D*. cf *indicum*: estimate = −0.175, *T. rugatulus*: estimate = −0.482, *C. formosanus*: estimate = −0.332, *R. speratus*: estimate = −0.147).

**Figure S4.**
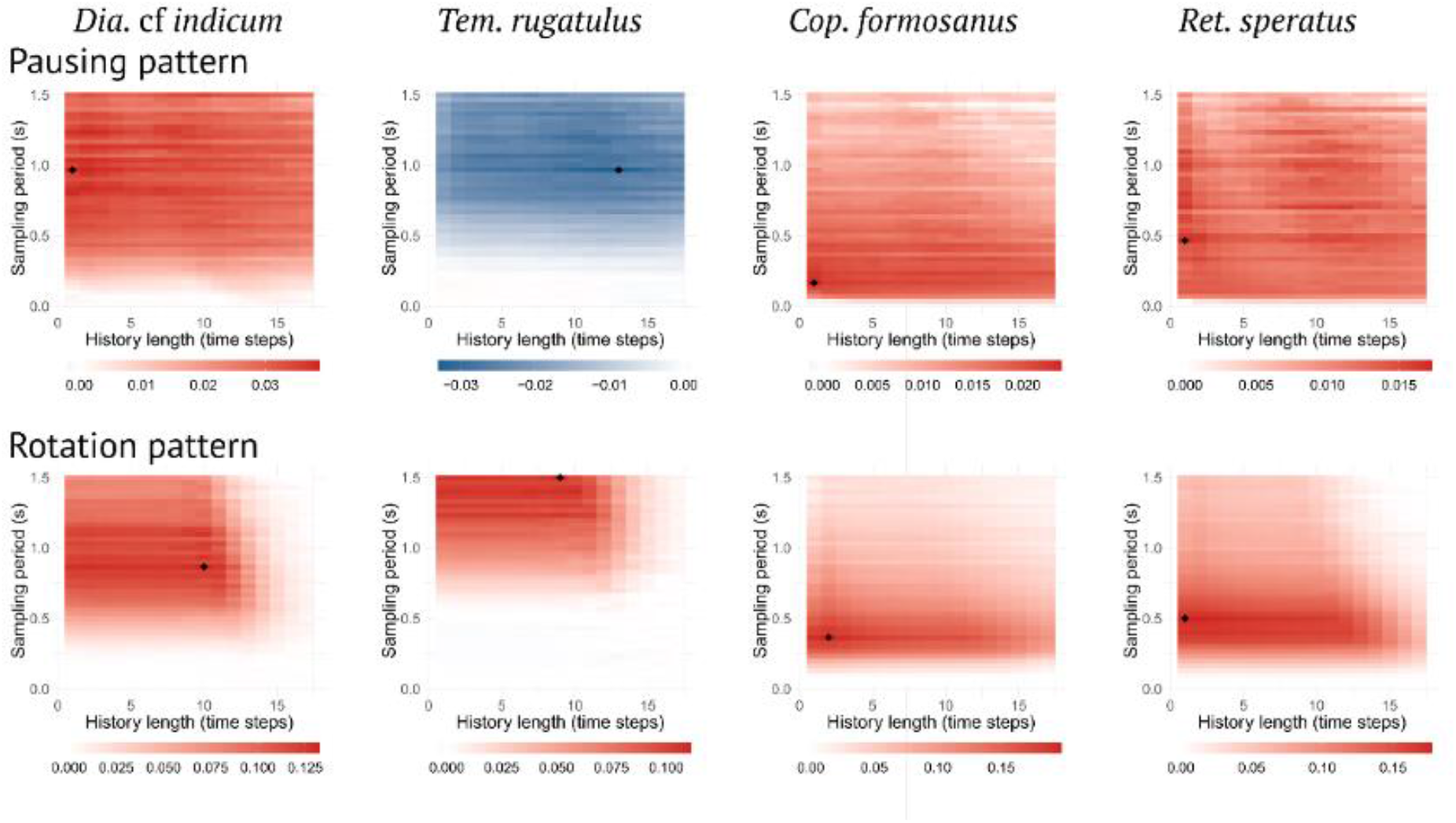
Landscape of net information transfer. Net transfer entropy (bits) as a function of the sampling period and of the history length. Colors indicate the intensity and predominant direction of information transfer (red, from leader to follower; blue, from follower to leader); the black diamond symbol indicates the configuration with maximum magnitude.

**Figure S5.**
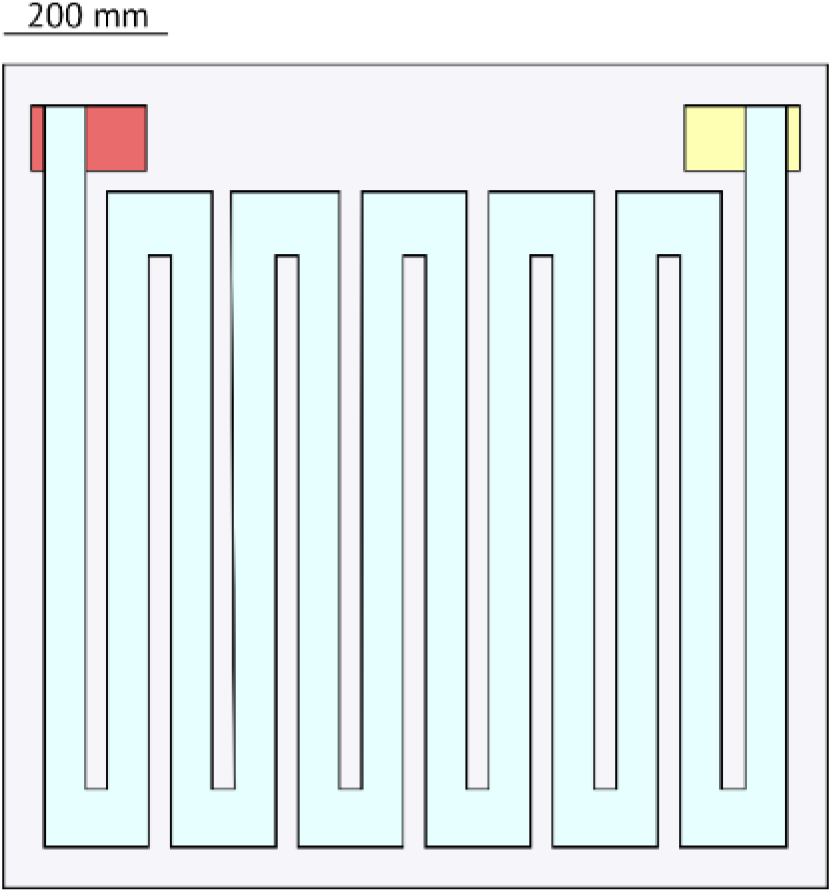
Experimental setup for the ant *Diacamma* cf *indicum*. The yellow box is the old nest, the red box is the new nest, and the light blue path indicates the bridge.

**Figure S6.**
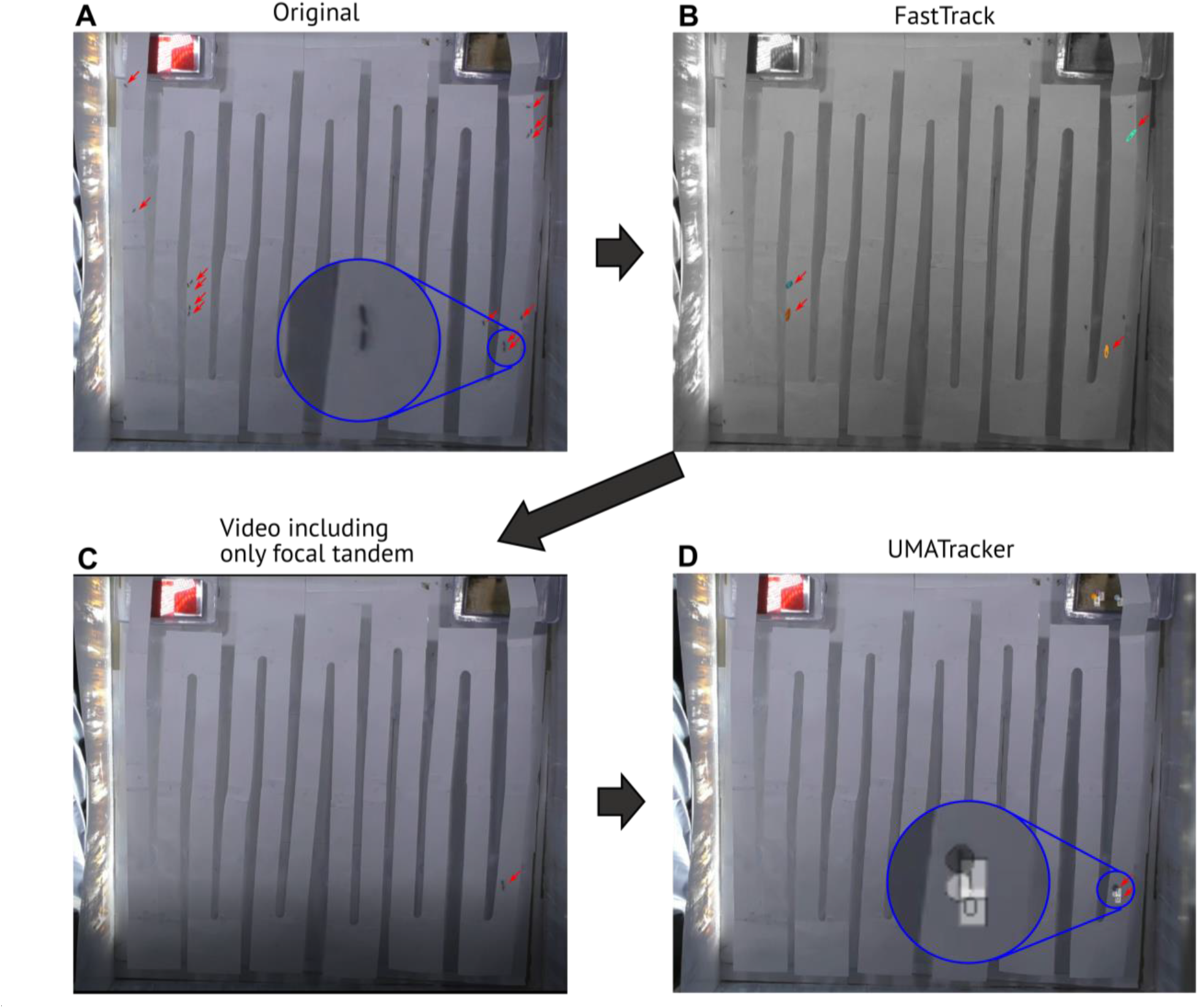
Procedure of movement tracking in this study. (A) In the original video, the total number of ants in a frame will vary according to time. There are 13 ant individuals in this frame, but we want to obtain coordinates of a focus tandem running pair in a blue circle. (B) FastTrack (Gallois, 2021) can handle the variable number of objects but cannot distinguish tandem partners. Here it detected four tandem pairs in the frame. (C) Using the rough coordinates of the focal tandem running pair, we produced a video masking other areas than the surrounding of the focal pair. There are usually only two focal individuals in this video, which is compatible with UMATracker (Yamanaka and Takeuchi, 2018). (D) UMATracker can get coordinates of the leader and follower separately during a tandem run.

**Figure S7.**
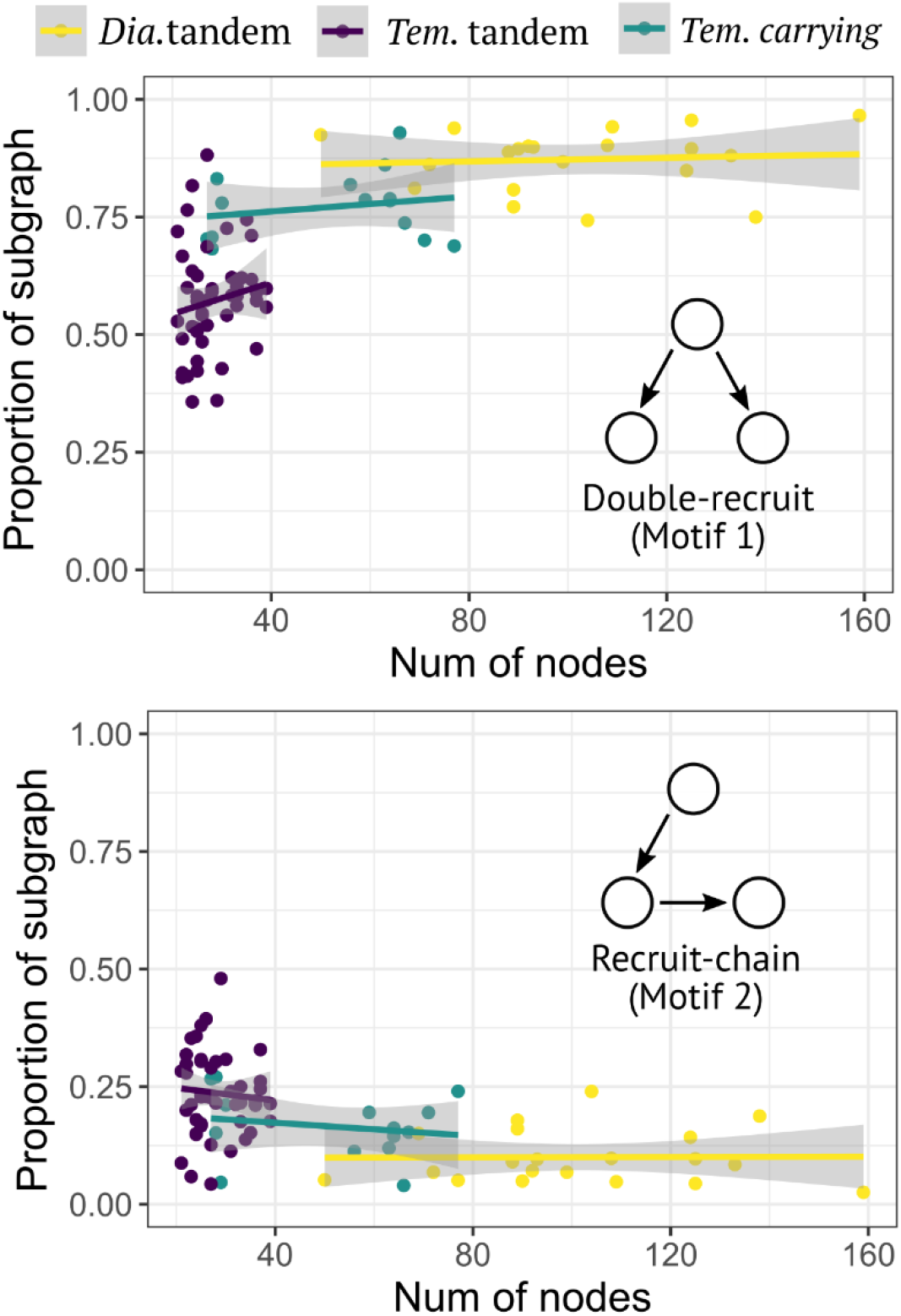
The relationship between the number of nodes and the proportion of each subgraph. All regression lines were not significant (liner model, P > 0.28). Networks with < 20 nodes were removed from the analysis.

**Figure S8.**
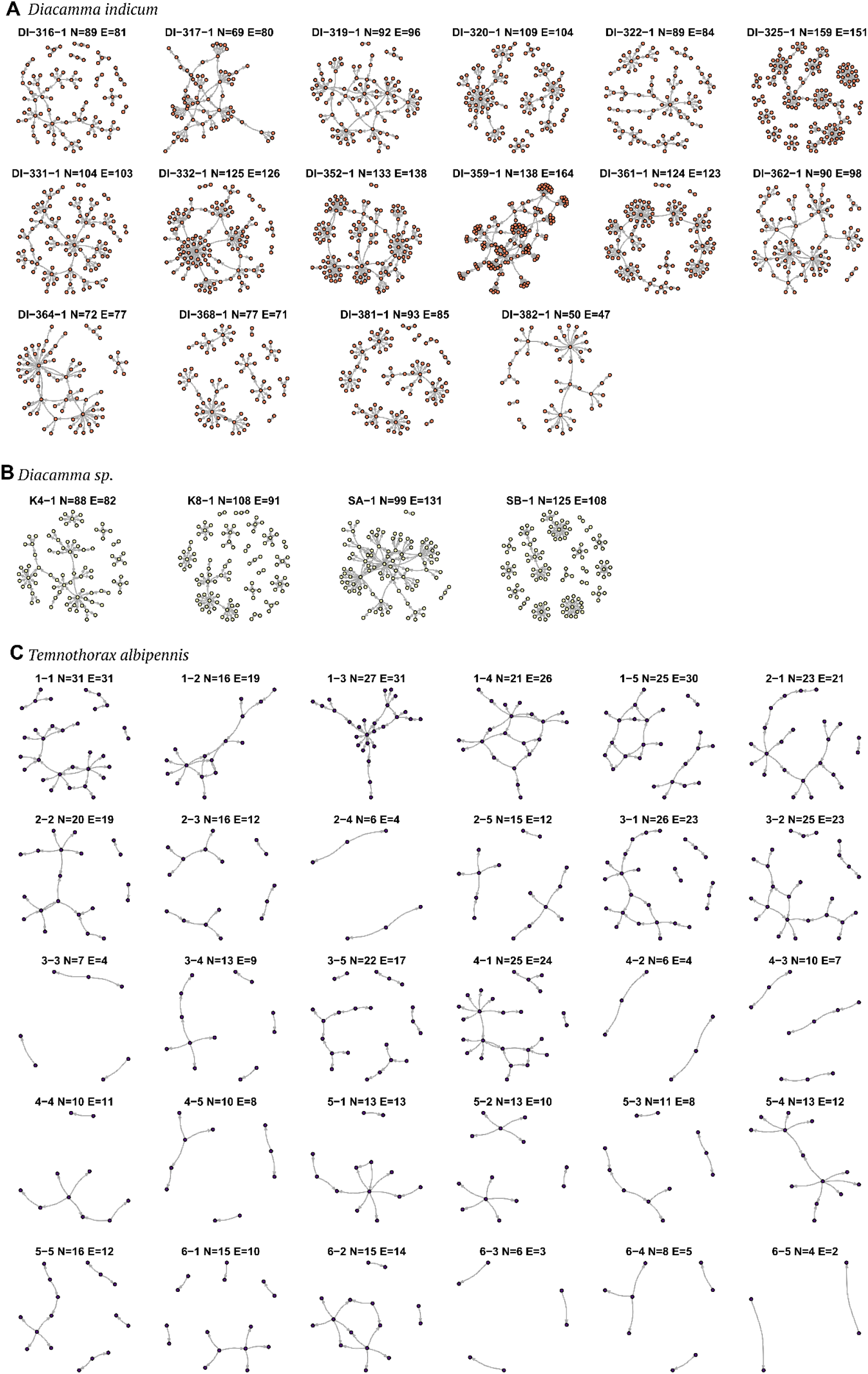
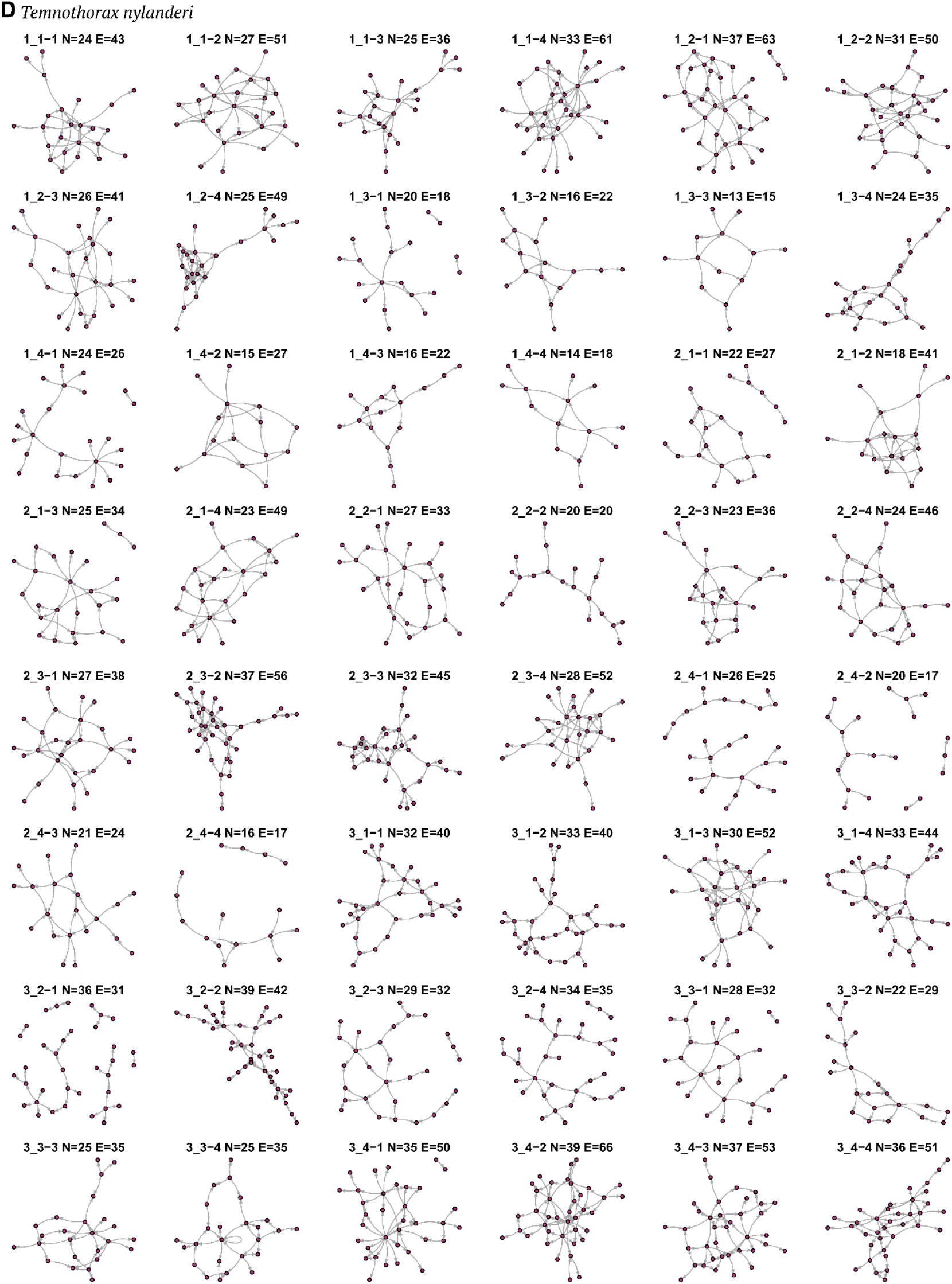
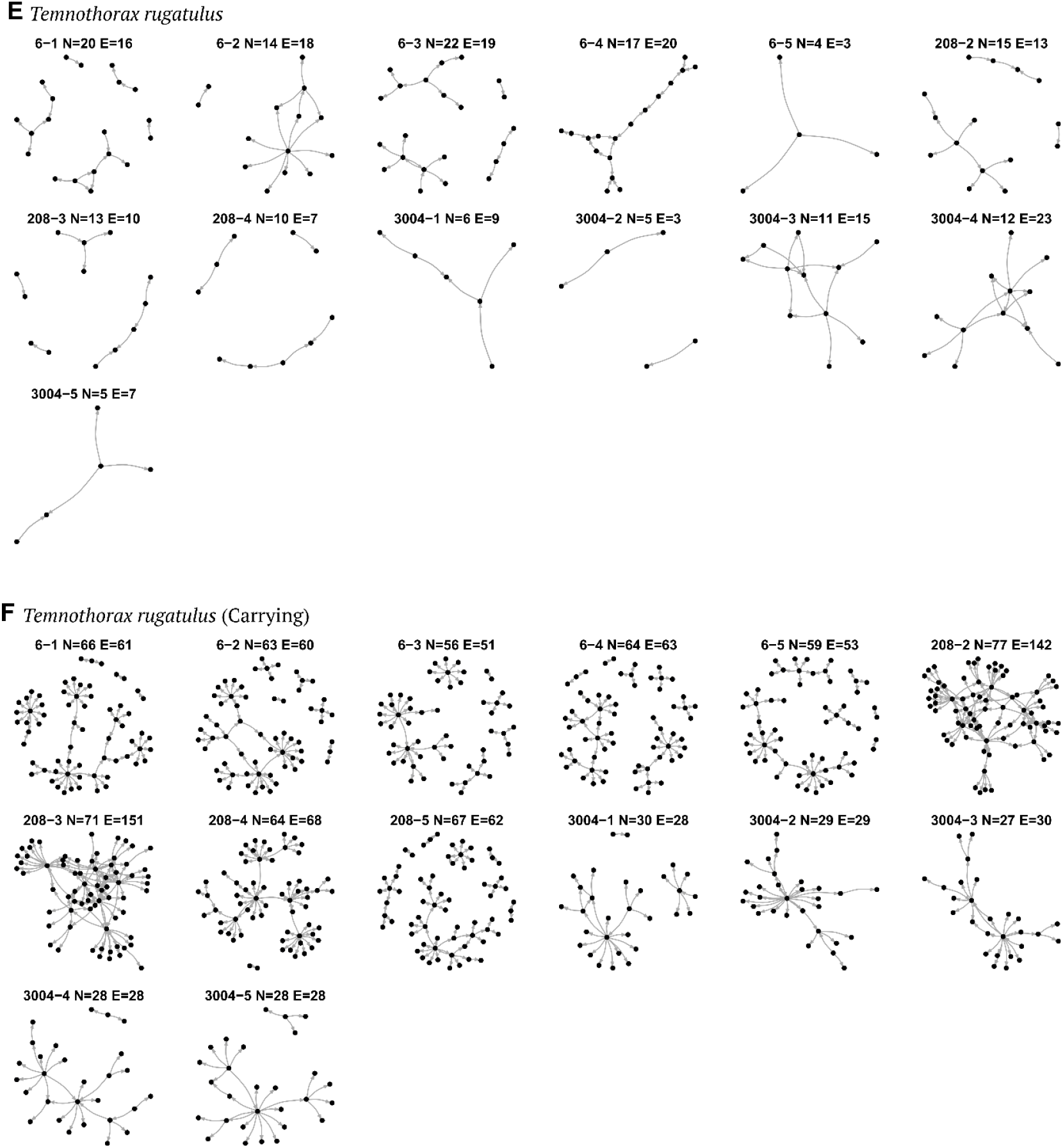
Visualization of all network structures used in this study

## References

Alleman A, Stoldt M, Feldmeyer B, Foitzik S. 2019. Tandem-running and scouting behaviour are characterized by up-regulation of learning and memory formation genes within the ant brain. Mol Ecol 28:2342–2359. doi:10.1111/mec.15079

Anoop K, Sumana A. 2015. Response to a change in the target nest during ant relocation. J Exp Biol 218:887–892. doi:10.1242/jeb.115246

Bates D, Mächler M, Bolker BM, Walker SC. 2015. Fitting linear mixed-effects models using lme4. J Stat Softw 67. doi:10.18637/jss.v067.i01

Bhattacharyya K, Kolay S, Annagiri S. 2021. The structure and importance of nest mounds in a tropical ant Diacamma indicum. Ecol Entomol 46:1324–1332. doi:10.1111/een.13079

Borowiec ML. 2019. Convergent evolution of the army ant syndrome and congruence in big-data phylogenetics. Syst Biol 68:642–656. doi:10.1093/sysbio/syy088

Buschinger A, Ehrhardt W, Winter U. 1980. The organization of slave raids in dulotic ants — a comparative study (Hymenoptera; Formicidae). Z Tierpsychol 53:245–264. doi:10.1111/j.1439-0310.1980.tb01053.x

Camazine S, Deneubourg J-L, Franks NR, Sneyd J, Theraulaz G, Bonabeau E. 2001. Self-organization in Biological Systems. Princeton: NJ: Princeton University Press.

Csardi G, Nepusz T. 2006. The igraph software package for complex network research. InterJournal Complex Syst 1695:1–9.

Donoghue MJ. 2005. Key innovations, convergence, and success: macroevolutionary lessons from plant phylogeny. Paleobiology 31:77–93. doi:10.1666/0094-8373(2005)031[0077:kicasm]2.0.co;2

Dornhaus A, Franks NR, Hawkins R, Shere HNS. 2004. Ants move to improve: Colonies of Leptothorax albipennis emigrate whenever they find a superior nest site. Anim Behav 67:959–963. doi:10.1016/j.anbehav.2003.09.004

Dussutour A, Simpson SJ. 2008. Description of a simple synthetic diet for studying nutritional responses in ants. Insectes Soc 55:329–333. doi:10.1007/s00040-008-1008-3

Foitzik S, Heinze J. 1998. Nest site limitation and colony takeover in the ant Leptothorax nylanderi. Behav Ecol 9:367–375. doi:10.1093/beheco/9.4.367

Franklin EL. 2014. The journey of tandem running: The twists, turns and what we have learned. Insectes Soc. doi:10.1007/s00040-013-0325-3

Franks Nigel R, Dornhaus A, Fitzsimmons JP, Stevens M. 2003. Speed versus accuracy in collective decision making. Proc R Soc London B 270:2457–2463. doi:10.1098/rspb.2003.2527

Franks Nigel R., Mallon EB, Bray HE, Hamilton MJ, Mischler TC. 2003. Strategies for choosing between alternatives with different attributes: Exemplified by house-hunting ants. Anim Behav 65:215–223. doi:10.1006/anbe.2002.2032

Franks NR, Richardson TO. 2006. Teaching in tandem-running ants. Nature 439:153. doi:10.1038/439153a

Fujiwara-Tsujii N, Tokunaga K, Akino T, Tsuji K, Yamaoka R. 2012. Identification of the tandem running pheromone in Diacamma sp. from Japan (Hymenoptera, Formicidae). Sociobiology 59:1281–1296.

Fukumoto Y, Abe T. 1983. Social organization of colony movement in the tropical ponerine ant, Diacamma rugosum (Le Guillou). J Ethol 1:101–108. doi:10.1007/BF02347836

Gallant JR, O’Connell LA. 2020. Studying convergent evolution to relate genotype to behavioral phenotype. J Exp Biol 223. doi:10.1242/jeb.213447

Gallois B. 2021. FastTrack: An open-source software for tracking varying numbers of deformable objects 1–19. doi:10.1371/journal.pcbi.1008697

Glaser SM, Feitosa RM, Koch A, Goß N, do Nascimento FS, Grüter C. 2021. Tandem communication improves ant foraging success in a highly competitive tropical habitat. Insectes Soc 68:161–172. doi:10.1007/s00040-021-00810-y

Glaser SM, Grüter C. 2022. Ancestral state reconstruction suggests repeated losses of recruitment communication during ant evolution (Hymenoptera: Formicidae). bioRxiv. doi:10.1101/2022.05.18.492496

Herbers JM. 1986. Nest site limitation and facultative polygyny in the ant Leptothorax longispinosus. Behav Ecol Sociobiol 19:115–122. doi:10.1007/BF00299946

Hothorn T, Bretz F, Westfall P. 2008. Simultaneous inference in general parametric models. Biometrical J. doi:10.1002/bimj.200810425

Jaekel M, Wake DB. 2007. Developmental processes underlying the evolution of a derived foot morphology in salamanders. Proc Natl Acad Sci U S A 104:20437–20442. doi:10.1073/pnas.0710216105

Kaur R, Anoop K, Sumana A. 2012. Leaders follow leaders to reunite the colony: Relocation dynamics of an Indian queenless ant in its natural habitat. Anim Behav 83:1345–1353. doi:10.1016/j.anbehav.2012.02.022

Kaur R, Joseph J, Anoop K, Sumana A. 2017. Characterization of recruitment through tandem running in an indian queenless ant Diacamma indicum. R Soc Open Sci 4. doi:10.1098/rsos.160476

Kolay S, Annagiri S. 2015a. Dual response to nest flooding during monsoon in an Indian ant. Sci Rep 5:1–9. doi:10.1038/srep13716

Kolay S, Annagiri S. 2015b. Tight knit under stress: Colony resilience to the loss of tandem leaders during relocation in an Indian ant. R Soc Open Sci 2. doi:10.1098/rsos.150104

Lautenschlager S, Figueirido B, Cashmore DD, Bendel EM, Stubbs TL. 2020. Morphological convergence obscures functional diversity in sabre-toothed carnivores: Sabre-tooth functional morphology. Proc R Soc B Biol Sci 287. doi:10.1098/rspb.2020.1818

Leonhardt SD, Menzel F, Nehring V, Schmitt T. 2016. Ecology and evolution of communication in social insects. Cell 164:1277–1287. doi:10.1016/j.cell.2016.01.035

Lizier JT, Prokopenko M. 2010. Differentiating information transfer and causal effect. Eur Phys J B 73:605–615. doi:10.1140/epjb/e2010-00034-5

Losos JB. 2011. Convergence, adaptation, and constraint. Evolution (N Y) 65:1827–1840. doi:10.1111/j.1558-5646.2011.01289.x

Menzel R, Brandt R, Gumbert A, Komischke B, Kunze J. 2000. Two spatial memories for honeybee navigation. Proc R Soc B Biol Sci 267:961–968. doi:10.1098/rspb.2000.1097

Meurville M-P, LeBoeuf AC. 2021. Trophallaxis: the functions and evolution of social fluid exchange in ant colonies (Hymeno ptera: Formicidae). Myrmecological News 31:1–30. doi:10.25849/myrmecol.news_031:00113

Milo R, Shen-Orr S, Itzkovitz S, Kashtan N, Chklovskii D, Alon U. 2002. Network motifs: simple building blocks of complex networks. Science (80-) 298:824–827.

Mizumoto N, Bourguignon T, Bailey NW. 2022. Ancestral sex-role plasticity facilitates the evolution of same-sex sexual behaviour. bioRxiv. doi:10.1101/2022.06.20.496918

Mizumoto N, Dobata S. 2019. Adaptive switch to sexually dimorphic movements by partner-seeking termites. Sci Adv 5:eaau6108. doi:10.1126/sciadv.aau6108

Mizumoto N, Lee S Bin, Valentini G, Chouvenc T, Pratt SC. 2021. Coordination of movement via complementary interactions of leaders and followers in termite mating pairs. Proc R Soc B Biol Sci 288:20210998. doi:10.1098/rspb.2021.0998

Mizumoto N, Rizo A, Pratt SC, Chouvenc T. 2020. Termite males enhance mating encounters by changing speed according to density. J Anim Ecol 89:2542–2552. doi:10.1111/1365-2656.13320

Möglich M. 1978. Social organization of nest emigration in Leptothorax (Hym., Form.). Insectes Soc 25:205–225. doi:10.1007/BF02224742

Moglich M, Maschwitz U, Holldobler B. 1974. Tandem calling: A new kind of signal in ant communication. Science (80-) 186:1046–1047.

Moore DG, Valentini G, Walker SI, Levin M. 2018. Inform: Efficient information-theoretic analysis of collective behaviors. Front Robot AI 5:1–14. doi:10.3389/frobt.2018.00060

Nelsen Matthew P., Ree RH, Moreau CS. 2018. Ant–plant interactions evolved through increasing interdependence. Proc Natl Acad Sci U S A 115:12253–12258. doi:10.1073/pnas.1719794115

Nelsen Matthew P, Ree RH, Moreau CS. 2018. Ant–plant interactions evolved through increasing interdependence. Proc Natl Acad Sci 115. doi:10.1073/pnas.1719794115

Perna A, Theraulaz G. 2017. When social behaviour is moulded in clay: on growth and form of social insect nests. J Exp Biol 220:83–91. doi:10.1242/jeb.143347

Pfeffer SE, Wittlinger M. 2016. Optic flow odometry operates independently of stride integration in carried ants. Science (80-) 353:1155–1157. doi:10.1126/science.aaf9754

Porfiri M. 2018. Inferring causal relationships in zebrafish-robot interactions through transfer entropy: a small lure to catch a big fish. Anim Behav Cogn 5:341–367. doi:10.26451/abc.05.04.03.2018

Pratt SC. 2005. Behavioral mechanisms of collective nest-site choice by the ant Temnothorax curvispinosus. Insectes Soc 52:383–392. doi:10.1007/s00040-005-0823-z

Pratt SC, Sumpter DJT, Mallon EB, Franks NR. 2005. An agent-based model of collective nest choice by the ant Temnothorax albipennis. Anim Behav 70:1023–1036. doi:10.1016/j.anbehav.2005.01.022

R Core Team. 2020. R: A language and environment for statistical computing.

Reeves DD, Moreau CS. 2019. The evolution of foraging behavior in ants (Hymenoptera: Formicidae). Arthropod Syst Phylogeny 77:351–363. doi:10.26049/ASP77-2-2019-10

Revell LJ. 2012. phytools: An R package for phylogenetic comparative biology (and other things). Methods Ecol Evol 3:217–223. doi:10.1111/j.2041-210X.2011.00169.x

Richardson TO, Coti A, Stroeymeyt N, Keller L. 2021. Leadership – not followership – determines performance in ant teams. Commun Biol 4:1–9. doi:10.1038/s42003-021-02048-7

Richardson TO, Mullon C, Marshall JAR, Franks NR, Schlegel T. 2018. The influence of the few: A stable ‘oligarchy’ controls information flow in house-hunting ants. Proc R Soc B Biol Sci 285. doi:10.1098/rspb.2017.2726

Sahu PK, Kolay S, Annagiri S. 2019. To reunite or not: A study of artificially fragmented Diacamma indicum ant colonies. Behav Processes 158:4–10. doi:10.1016/j.beproc.2018.10.017

Sasaki T, Colling B, Sonnenschein A, Boggess MM, Pratt SC. 2015. Flexibility of collective decision making during house hunting in Temnothorax ants. Behav Ecol Sociobiol 69:707–714. doi:10.1007/s00265-015-1882-4

Sasaki T, Danczak L, Thompson B, Morshed T, Pratt SC. 2020. Route learning during tandem running in the rock ant Temnothorax albipennis. J Exp Biol 223:jeb221408. doi:10.1242/jeb.221408

Schielzeth H, Dingemanse NJ, Nakagawa S, Westneat DF, Allegue H, Teplitsky C, Réale D, Dochtermann NA, Garamszegi LZ, Araya-Ajoy YG. 2020. Robustness of linear mixed-effects models to violations of distributional assumptions. Methods Ecol Evol 2041–210X.13434. doi:10.1111/2041-210x.13434

Schneider CA, Rasband WS, Eliceiri KW, Instrumentation C. 2017. NIH Image to ImageJ: 25 years of Image Analysis 9:671–675.

Schreiber T. 2000. Measuring information transfer. Phys Rev Lett 85:461–464. doi:10.1103/PhysRevLett.85.461

Schultheiss P, Raderschall CA, Narendra A. 2015. Follower ants in a tandem pair are not always naïve. Sci Rep 5. doi:10.1038/srep10747

Shimoji H, Dobata S. 2022. The build-up of dominance hierarchies in eusocial insects. Philos Trans R Soc Lond B Biol Sci 377:20200437. doi:10.1098/rstb.2020.0437

Silva JP, Valadares L, Vieira MEL, Teseo S, Châline N. 2021. Tandem running by foraging Pachycondyla striata workers in field conditions vary in response to food type, food distance, and environmental conditions. Curr Zool 67:541–549. doi:10.1093/cz/zoab050

Stroeymeyt N, Giurfa M, Franks NR. 2010. Improving decision speed, accuracy and group cohesion through early information gathering in house-hunting ants. PLoS One 5. doi:10.1371/journal.pone.0013059

Stroeymeyt N, Robinson EJH, Hogan PM, Marshall JAR, Giurfa M, Franks NR. 2011. Experience-dependent flexibility in collective decision making by house-hunting ants. Behav Ecol 22:535–542. doi:10.1093/beheco/arr007

Theraulaz G, Bonabeau E. 1995. Coordination in distributed building. Science (80-) 269:686–688. doi:10.1126/science.269.5224.686

Therneau TM. 2015. coxme: mixed effects Cox models.

Tsuji K. 2021. Reproductive differentiation and conflicts in Diacamma: A model system for integrative sociobiology. Asian Myrmecology 13. doi:10.20362/am.013007

Tsuji K, Egashira K, Holldobler B. 1999. Regulation of worker reproduction by direct physical contact in the ant Diacamma sp. from Japan. Anim Behav 58:1329. doi:10.1006/anbe.1999.1263

Valentini G, Masuda N, Shaffer Z, Hanson JR, Sasaki T, Walker SI, Pavlic TP, Pratt SC. 2020a. Division of labour promotes the spread of information in colony emigrations by the ant Temnothorax rugatulus. Proc R Soc B Biol Sci 287. doi:10.1098/rspb.2019.2950

Valentini G, Mizumoto N, Pratt SC, Pavlic TP, Walker SI. 2020b. Revealing the structure of information flows discriminates similar animal social behaviors. Elife 9:e55395. doi:10.7554/eLife.55395

Viginier B, Peeters C, Brazier L, Doums C. 2004. Very low genetic variability in the Indian queenless ant Diacamma indicum. Mol Ecol 13:2095–2100. doi:10.1111/j.1365-294X.2004.02201.x

Warton DI, Hui FKC. 2011. The arcsine is asinine: The analysis of proportions in ecology. Ecology 92:3–10. doi:10.1890/10-0340.1

Waters JS, Fewell JH. 2012. Information processing in social insect networks. PLoS One 7. doi:10.1371/journal.pone.0040337

Wilson EO. 1971. The Insect Societies. Cambridge: Harvard University Press.

Win AT, Machida Y, Miyamoto Y, Dobata S, Tsuji K. 2018. Seasonal and temporal variations in colony-level foraging activity of a queenless ant, Diacamma sp., in Japan. J Ethol 36:277–282. doi:10.1007/s10164-018-0558-8

Wittlinger M, Wehner R, Wolf H. 2006. The ant odometer: Stepping on stilts and stumps. Science (80) 312:1965–1967. doi:10.1126/science.1126912

Yamanaka O, Takeuchi R. 2018. UMATracker: An intuitive image-based tracking platform. J Exp Biol 221:1–24. doi:10.1242/jeb.182469

